# Exercise enhances motor skill learning by neurotransmitter switching in the adult midbrain

**DOI:** 10.1101/613539

**Authors:** Hui-quan Li, Nicholas C. Spitzer

## Abstract

Physical exercise promotes motor skill learning in normal individuals and those with neurological disorders but its mechanism of action is unclear. We found that one week of voluntary wheel running enhances the acquisition of motor skills in adult mice. One week of running also induces switching from ACh to GABA expression in neurons in the caudal pedunculopontine nucleus (cPPN). The switching neurons make projections to the substantia nigra (SN), ventral tegmental area (VTA) and ventrolateral-ventromedial nuclei of the thalamus (VL-VM), which regulate acquisition of motor skills. Use of viral vectors to override transmitter switching blocks the beneficial effect of running on motor skill learning. We suggest that neurotransmitter switching provides the basis by which sustained running benefits motor skill learning, presenting a new target for clinical treatment of movement disorders.

## Main

Motor skill learning is fundamental to everyday life and is regulated by a neural network involving the cortex, thalamus, basal ganglia, brain stem, cerebellum and spinal cord (*1,2*). Both neuronal and glial plasticity are essential for motor skill learning and disruption of this plasticity causes motor deficits (*3,4*). Aerobic physical exercise promotes the ability to acquire new motor skills (*5*) and serves as a therapy for many motor disorders (*6–8*), but its basis of action is not well understood. Running is a natural motor activity for mice (*9*) and generates plasticity in multiple brain regions (*4,10,11*). Neurotransmitter switching is a newly appreciated form of plasticity that refers to the ability of neurons to change their transmitter identity in response to sustained stimuli, typically leading to changes in behavior (*12*). We hypothesized that chronic running induces neurotransmitter switching in a circuit that is important for motor skill learning.

## Results

### Sustained running enhances motor skill learning

We investigated the impact of chronic voluntary running on motor skill learning by exposing adult mice to running wheels for one week (Fig. 1a). Each mouse spent consistent time on the wheel every day, indicating continuous interest, and their running skill improved as measured by increased running speed, increased duration of running episodes and more stable movement on the wheel (Fig. 1b and Supplementary Fig. 1a,b). At the end of the running period we assessed performance on the rotarod and balance beam to evaluate motor skill acquisition. In comparison to mice without running wheels, mice that ran for one week demonstrated enhanced learning of motor skills, mastering an accelerating rotarod more rapidly and accommodating to balance beams more quickly (Fig. 1c-f and Supplementary Fig. 1c,d).

**Fig. 1.**
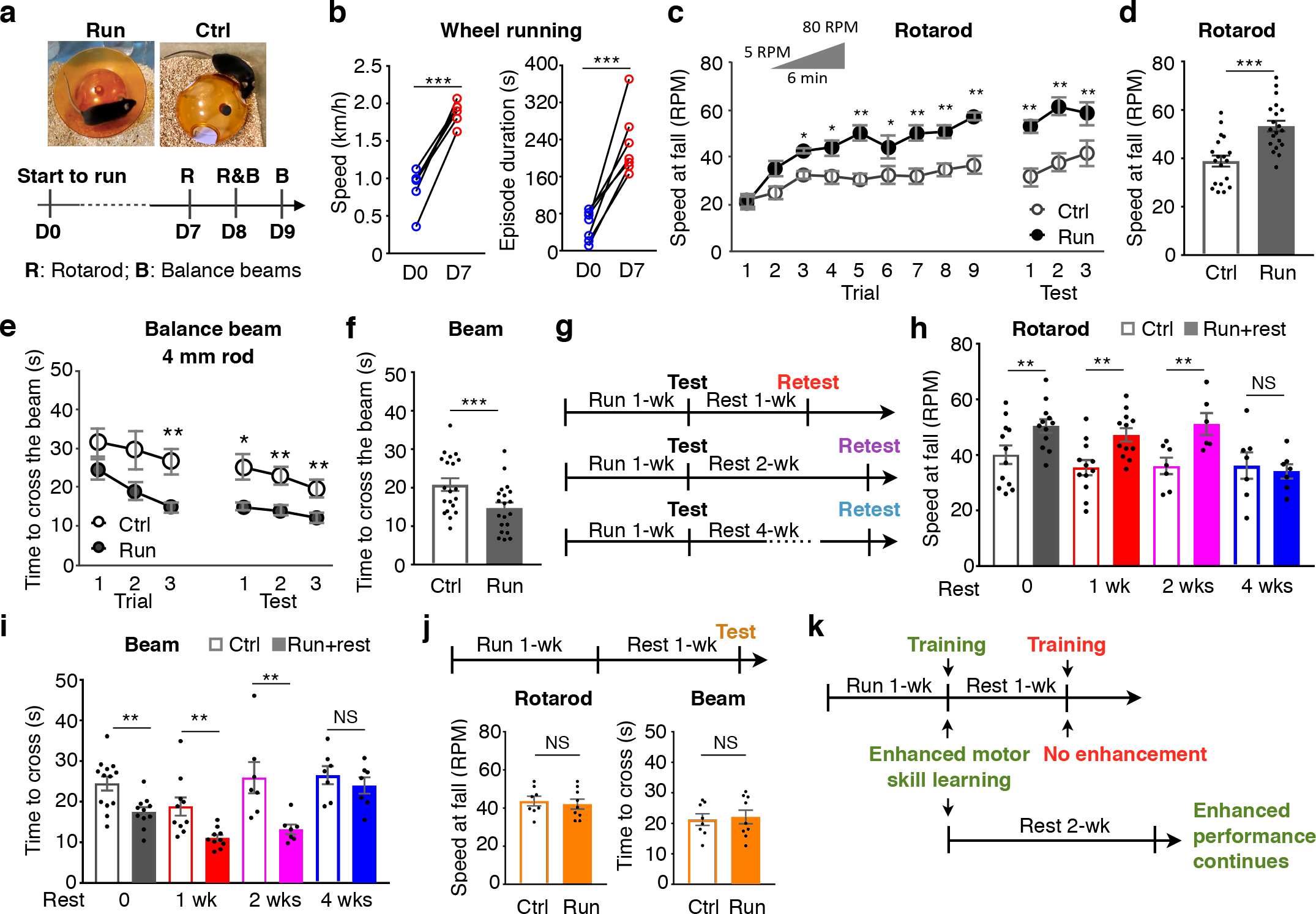
Sustained running enhances motor skill learning. **a,** Top: Runner mice were provided with running wheels in their home cage and control mice were housed with wheelbases. Bottom: the experimental timeline for running and behavioral tests. D, day. **b,** Mean speed (left, n=6 animals/group) and mean episode duration (right, n=7 animals/group) of mice running on day 0 (naive) and on day 7 (1-wk trained). **c**, Speed at fall on a rotarod in accelerative mode of each trial during training or of each test on the day after training. **d,** Mean speed at fall on a rotarod in accelerative mode in three tests on the day after training. **e**, Time to cross a 1 meter long, 0.75 meter high, 4 mm diameter rod balance beam during each trial of training or each test on the day after training. **f,** Mean time to cross the 4 mm rod beam in three tests on the day after training. For (**c**-**f**), n=19 for Ctrl and 20 for Run. **g,** Experimental design for immediate behavioral testing and retesting. **h,i**, Mice that had run for 1 week or non-runner controls were tested with (**h**) rotarod and (**i**) balance beam (4 mm rod). After rest for 1, 2 or 4 weeks, mice were retested. n=12 animals/group for 0 wk, 10 animals/group for 1 wk, 7 animals/group for 2 wks, and 7 animals/group for 4 wks. **j,** Top: experimental design for delayed behavioral testing. Bottom: performances on rotarod (left) and balance beam (right, 4 mm rod) after 1 week of running followed by 1 week of rest, compared to controls that never ran on a running wheel. n=8 animals for Ctrl and 9 animals for Run+Rest. **k,** Summary of time dependence of enhanced motor skill learning. Statistical significance \**p*<0.05, \*\**p*<0.01, \*\*\**p*<0.001 was assessed by paired t-test (**b**) or Welch’s t-test (other panels). NS, not significant. Data shown are mean±SEM.

We calculated the slopes of learning curves to assess the speed of motor skill learning. Mean data points for each training trial were plotted and fitted and the coefficient of determination (R^2^) was used to justify the fit. The mean speed at fall from a rotarod for the nine trials on the training day was fitted by a one-phase association model (R^2^>0.96 for controls; R^2^>0.95 for runners).

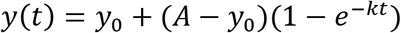

*y*(*t*) is the speed at fall on trial number *t*, *y*_0_ is the speed at fall for the first trial (*t* = 0), *A* is the plateau and *k* is a rate constant.

The slopes were calculated with

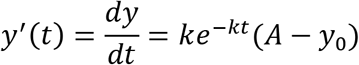

To calculate the initial slope, *y*’(0), we used

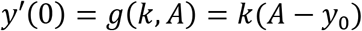

The standard errors were calculated with

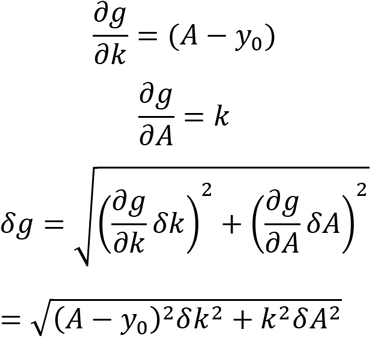

The values of the slopes are presented as *g* ± *δg* (mean ± sem). Initial slopes were 17±4 rpm/trial for runners and 5±2 rpm/trial for controls, significantly steeper for the runners (*p*=0.01, Welch’s t-test) (Fig. 1c).

The mean time to cross a 4 mm rod beam for the three trials on the training day was fitted by linear regression (R^2^>0.98 for controls; R^2^>0.99 for runners)

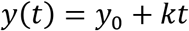

*y*(*t*) is the time to cross the beam on trial number t, *y*_0_ is the time for the first trial (*t* = 0) and *k* is the slope

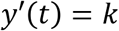

The values of the slopes are presented as *k* ± *δk* (mean ± sem). The slopes were −4.7 ± 0.5 s/trial for runners and −2.4 ± 0.3 s/trial for controls, again significantly steeper for runners (*p*<0.001, Welch’s t-test) (Fig. 1e). Additionally, sustained running improved rotorod and balance beam test performance (Fig. 1d,f and Supplementary Fig. 1d) but did not affect basal locomotor activity as measured by infrared beam crossings in home cages (Supplementary Fig. 1e,f).

To explore further the effect of running on enhancement of motor skill learning, we removed running wheels from mouse cages after the first motor skill tests and retested the mice after different resting periods (Fig. 1g). Enhanced performance on both rotarod and balance beam was sustained up to 2 weeks but not 4 weeks (Fig. 1h,i). Moreover, when mice were not trained on the rotarod and balance beam until 1 week after running, their motor skill learning was not enhanced (Fig. 1j). This result suggests that running creates a sensitive period in the adult brain during which motor skill learning is improved and shows that the gain in motor skills persists longer than the duration of this period (Fig. 1k).

### Running induces transmitter switching in the midbrain cPPN

Because transmitter switching is activity-dependent (*13–15*), we first searched for c-fos expression (Fig. 2a) to determine sites of increased brain activity associated with running and to identify neurons likely to switch their transmitter. Mice that ran for 1 week exhibited a 6-fold increase in the number of c-fos+ neurons in the pedunculopontine nucleus (PPN) of the midbrain compared to non-runner controls when examined immediately after the end of the last running period (Fig. 2b,c). The dentate gyrus also showed increased c-fos expression in runners, as expected (*16*). The PPN was an attractive candidate for running-induced transmitter switching because it regulates gait and balance control in health (*17,18*) and disease (*19*). However, the rostral and caudal PPN (rPPN and cPPN) are distinct. Both contain glutamatergic, GABAergic and cholinergic neurons, but there are important differences in the proportions of their transmitter phenotypes (*20*), their projection targets and activity during movement (*18,21*), and their roles in behavior (*22–24*). Accordingly we looked for differences in the activation of the rPPN and cPPN.

**Fig. 2.**
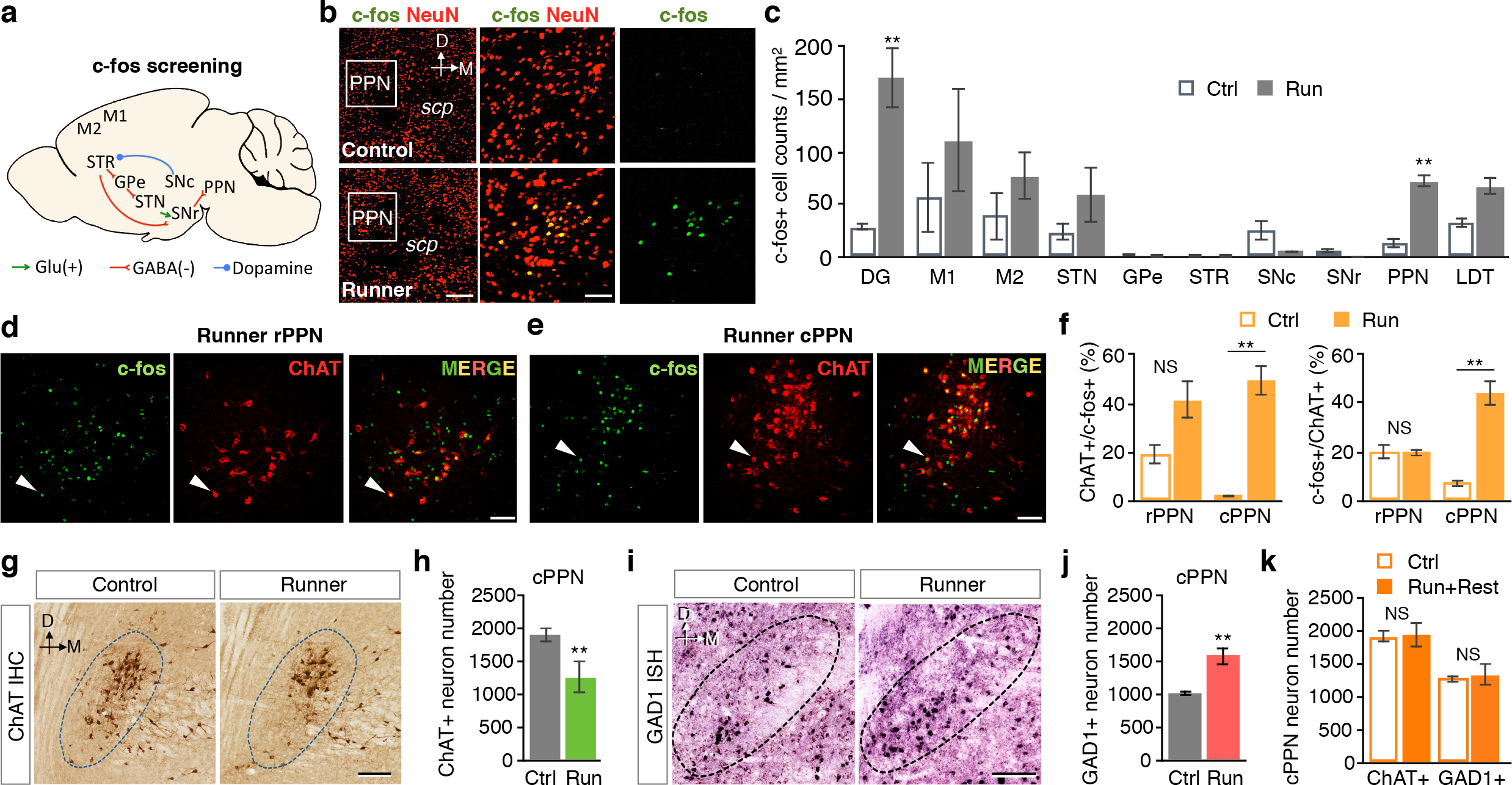
Running activates the pedunculopontine nucleus (PPN) and triggers neurotransmitter switching in the caudal PPN. **a,** Motor circuitry screened for activity. **b,** Double staining of neuronal marker NeuN and activity marker c-fos in a control (upper) and 1-week runner (lower). Middle and right panels are higher magnifications of the white boxes in the left panels. *Scp*, superior cerebellar peduncle. Dorsal (D) and medial (M) axes are shown at top. Scale bar, (left) 200 μm, (middle) 50 μm. **c,** c-fos immunoreactive neuron number in each region or nucleus of controls and 1-week runner mice. DG, dentate gyrus; M1, primary motor cortex; M2, secondary motor cortex; STN, subthalamic nucleus; GPe, globus pallidus external; STR, striatum; SNc, substantia nigra *pars* compacta; SNr, substantia nigra *pars* reticulata; PPN, pedunculopontine nucleus; LDT, laterodorsal tegmental nucleus. n=12 sections from 4 animals/group. **d,e,** Double staining of ChAT and c-fos in the rostral (**d**) or caudal (**e**) PPN of a 1-wk runner. White arrowheads identify examples of c-fos+ChAT+ cells. Scale bar, 100 μm. **f,** Percentage of the c-fos+ChAT+ neurons in the total ChAT+ neurons (left, *p*=0.07 for the rPPN) or in the total c-fos+ neurons (right). n=3 animals/group. **g,** 3,3’-diaminobenzidine (DAB) staining of ChAT in the cPPNs of a control and 1-week runner. Dotted lines outline the PPN. Dark brown stain identifies ChAT+ neurons. Scale bar, 100 μm. **h,** Stereological counts of (**g**). n=6 animals/group. **i,** *In situ* hybridization of GAD1 in the cPPN of a control and 1-week runner. Scale bar, 100 μm. **j,** Stereological counts of (**i**). n=7 animals/group. **k,** Stereological counts of ChAT+ or GAD1+ neurons in the cPPN of mice that experienced 1 week of running followed by 1 week of rest and of control mice that never ran on a running wheel. n=5 animals/group. Statistical significance \*\**p*<0.01 was assessed by Mann–Whitney U test. NS, not significant. Data shown are mean± SEM.

Specifically, we asked whether running activates cholinergic neurons differently in the rPPN and cPPN, because cholinergic neurons are involved in the gait and postural disorders of Parkinson’s disease (*25*). One week of running increased the number of c-fos+ neurons in both rPPN and cPPN (Supplementary Fig. 2). However, running dramatically increased the percentage of choline acetyltransferase (ChAT)+c-fos+ neurons in the ChAT+ neuron population by 22-fold in the cPPN but increased the percentage by only 2.1-fold in the rPPN (Fig. 2d,e, the left panel of 2f, and Supplementary Fig. 3a). Moreover, the percentage of ChAT+c-fos+ neurons in the c-fos+ neuron population was not different between runners and controls in the rPPN but increased 6-fold between runners and controls in the cPPN (Fig. 2f, right). Significantly, running increased the number of non-cholinergic cfos+ neurons in the cPPN by only 1.5-fold, much less than the 15-fold increase in the number of ChAT+c-fos+ neurons (Supplementary Fig. 3a). The increased proportion of c-fos+ neurons in the population of ChAT+ cPPN neurons and the increased proportion of ChAT+ neurons in the population of c-fos+ cPPN neurons occurred after as early as 3 days of running (Supplementary Fig. 3b,c). These results show that cPPN cholinergic neurons are more strongly activated by sustained running than rPPN cholinergic neurons or cPPN non-cholinergic neurons, identifying cPPN cholinergic neurons as candidates for transmitter switching.

**Fig. 3.**
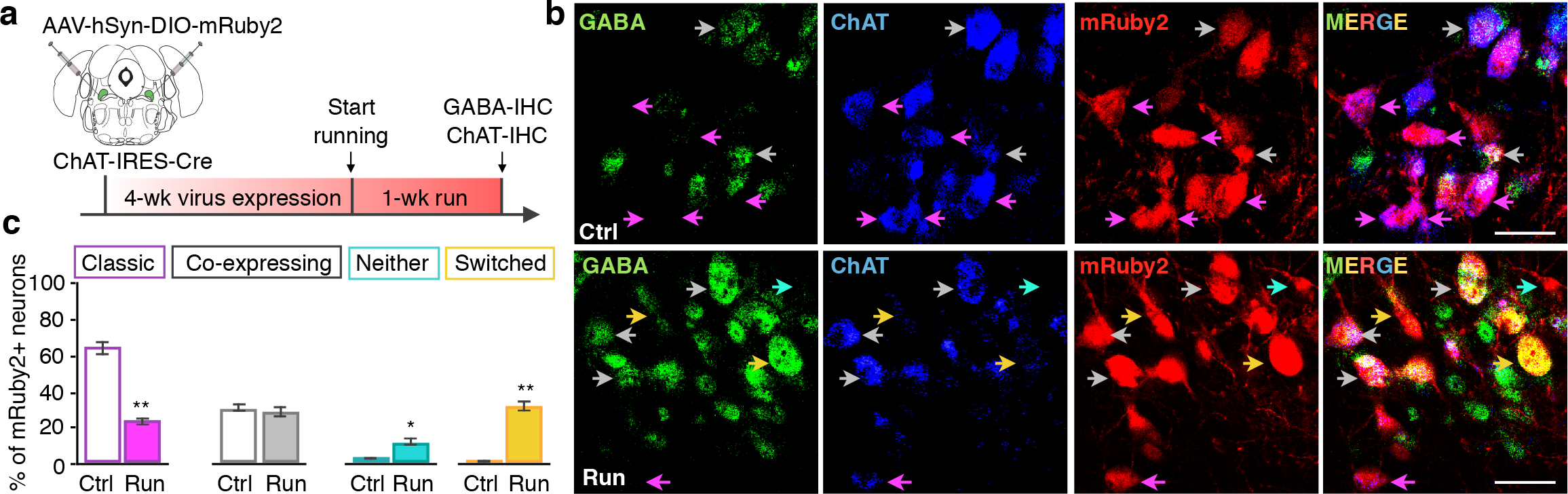
Cholinergic cPPN neurons lose ChAT and gain GABA. **a,** Experimental strategy to permanently label cholinergic cPPN neurons with mRuby2 fluorescent protein by expressing Cre-dependent AAV-DIO-mRuby2 in ChAT-Cre mice and determine whether the loss of ChAT and gain of GABA occur in mRuby2+ neurons. **b,** Double immunostaining of GABA and ChAT in the cPPN of a control and a 1-week runner ChAT-Cre mouse expressing AAV-DIO-mRuby2 in the cPPN. Purple arrows, mRuby2 neurons that express ChAT but not GABA (classic ChAT neurons). Grey arrows, mRuby2 neurons that express both ChAT and GABA (co-expressing neurons). Yellow arrows, mRuby2 neurons that express GABA but not ChAT (switched neurons). Cyan arrow, an mRuby2 neuron that express neither GABA nor ChAT. Scale bar, 50 μm. **c,** The percentage of each class of mRuby2+ neurons (n= 844 for controls and 738 cells for runners). n= 3 animals/group.

Indeed, chronic running for one week was accompanied by a decrease in the number of cPPN neurons expressing both ChAT (Fig. 2g,h) and the vesicular acetylcholine transporter (VAChT) (Supplementary Fig. 4). This change was accompanied by an equal increase in the number of cPPN neurons expressing the gene encoding glutamic acid decarboxylase (GAD1) that generates GABA (Fig. 2i,j). No neurogenesis or apoptosis was observed in the cPPN of either control or runner mice (Supplementary Fig. 5). These results suggest that ~600 cPPN neurons switched their transmitter from ACh to GABA. There was no change in the number of ChAT+ neurons in the rPPN or the adjacent lateral dorsotegmental nucleus and no difference in the number of neurons expressing the vesicular glutamate transporter 2 (vGluT2) in the cPPN (Supplementary Fig. 6).

**Fig. 4.**
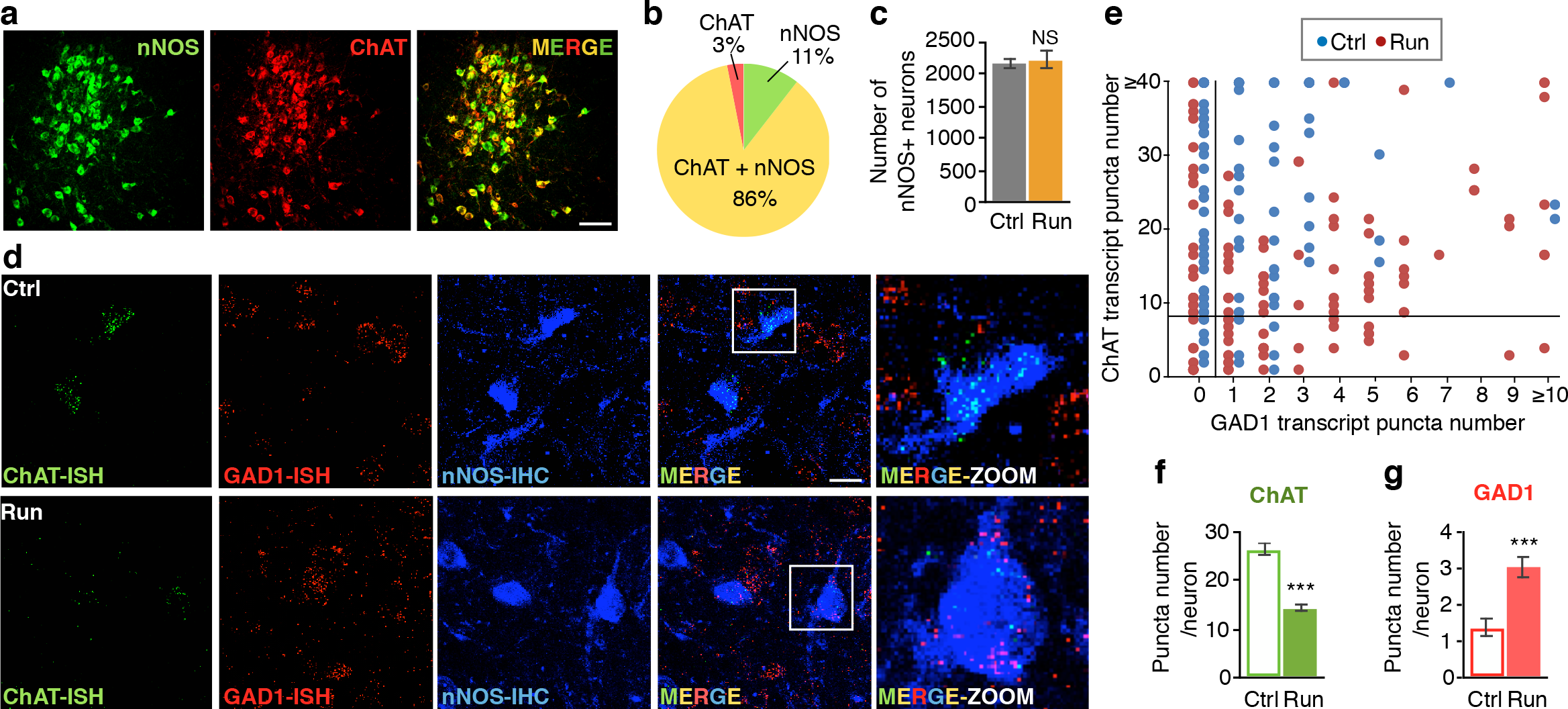
A subset of cPPN nNOS neurons of runner mice lose ChAT transcripts and gain GAD1 transcripts. **a,** Double staining of the cPPN in a non-runner control mouse for nNOS (green) and ChAT (red). Scale bar, 100 μm. **b,** Co-localization of nNOS and ChAT in the cPPN. n=1068 cells from 3 non-runner control mice. **c,** Stereological counts of DAB staining of nNOS in control and 1-week runner cPPNs. n=6 animals/group. Mann–Whitney U test. NS, not significant. **d,** Triple staining of ChAT and GAD1 transcripts and nNOS protein in the cPPN of a control and a 1-week runner mouse. Right panels are boxed regions in merged images at higher magnification. Scale bar, 20 μm. **e,** Scatterplot of numbers of *in situ* stained ChAT puncta (y-axis) against numbers of *in situ* stained GAD1 puncta (x-axis). Each dot represents one neuron. The vertical line divides neurons that contain zero from those containing more GAD1 transcript puncta and the horizontal line divides neurons that contain less from those containing more than eight ChAT transcript puncta. **f,g,** Y-axes are the mean number of ChAT (**f**) and GAD1 (**g**) fluorescent puncta in single nNOS+ neurons. For (**e**-**g**), n=4 animals/group; n=123 cells for Ctrl and 137 cells for Run. Statistical significance \**p*<0.05, \*\*\**p*<0.001 was assessed by Welch’s t-test. Data shown are mean± SEM.

**Fig. 5.**
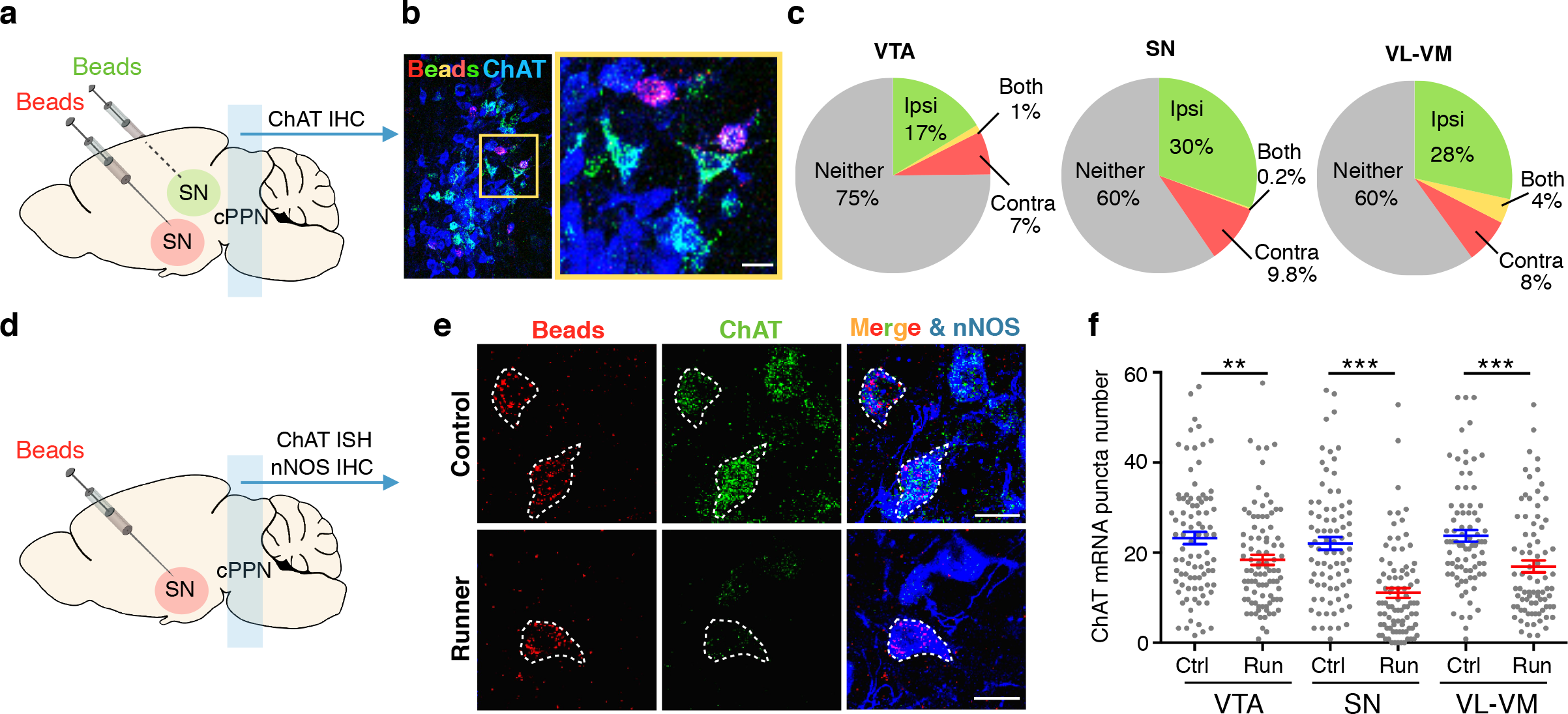
Running reduces the number of ChAT transcripts in a subset of cholinergic cPPN neurons that project to the VTA, SN and VL-VM. **a,** Experimental design to validate cholinergic innervation of target nuclei by the cPPN. Red or green retrobeads (beads) were injected bilaterally into target nuclei with one color per side. Substantia nigra (SN) is shown as an example. **b,** Triple-labeled retrobeads and ChAT in a coronal section of cPPN in a mouse injected with retrobeads. Scale bar, 50 μm. (**c**) Summary of the percentage of cholinergic neurons (ChAT+) that project to corresponding nuclei. Ipsi, ipsilaterally. Contra, contralaterally. Both, both ipsi-and contralaterally. Neither, no retrobeads. n=3 animals/examined region. n= 829 cells for the VTA, 806 cells for the SN, and 812 for the VL-VM. **d,** Experimental design to identify the target(s) of neurons with decreased numbers of ChAT transcripts. SN is shown as an example. **e,** Triple-labeling of retrobeads, ChAT mRNA transcripts, and nNOS proteins in both control and runner cPPNs. Scale bar, 20 μm. **f,** Y-axis is the mean number of ChAT fluorescent puncta in single nNOS+ cells. Each dot represents one neuron. n=4 animals/group. n=89 cells for VTA-Ctrl, 91 for VTA-Run, 81 for SN-Ctrl, 87 for SN-Run, 82 for VL-VM-Ctrl, 80 for VL-VM-Run. Statistical significance \*\**p*<0.01, \*\*\**p*<0.001 was assessed by Welch’s t-test. Data shown are mean± SEM.

**Fig. 6.**
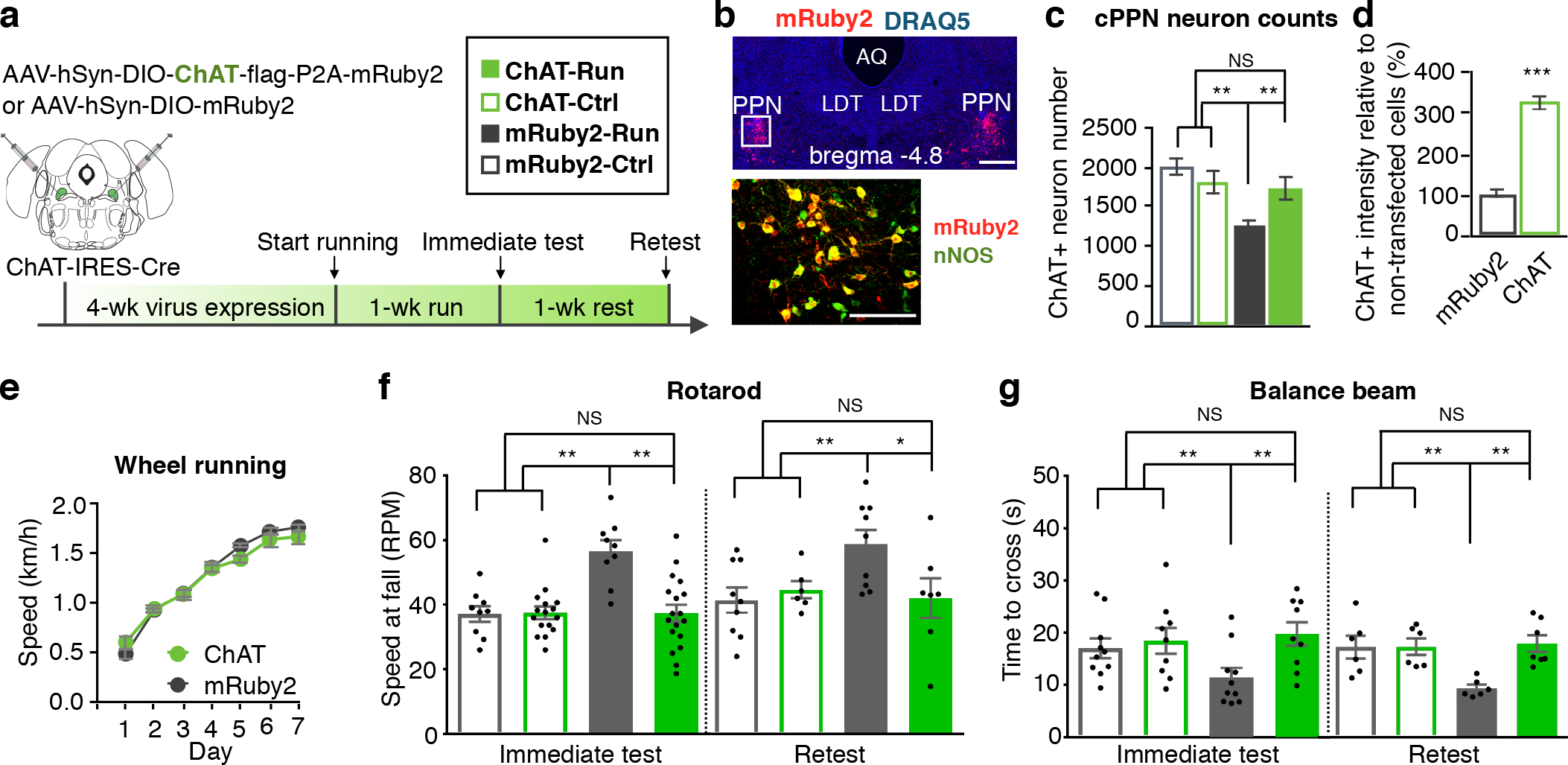
Loss of ChAT in cholinergic cPPN neurons is necessary for running-enhanced motor skill learning. **a,** Experimental design to override the loss of ChAT in ChAT-Cre neurons and determine behavioral relevance. **b,** Top: mRuby2 and nuclear marker DRAQ-5 show mRuby2 expression in a coronal cPPN section of an animal bilaterally-injected with AAV-DIO-ChAT-P2A-mRuby2 constructs. Coordinates adapted to Allen Brain Atlas. AQ, aqueduct. LDT, laterodorsal tegmental nucleus. Scale bar, 500 μm. Bottom: double-labeled image of nNOS and mRuby2 in boxed region of the upper image. Scale bar, 200 μm. **c,** ChAT+ neuron number in the cPPN of runners and non-running controls that were injected with AAV-DIO-mRuby2 or AAV-DIO-ChAT. n=6 animals/group. Nonparametric Kruskal–Wallis test followed by Dunn’s correction. **d,** ChAT fluorescence intensity for mRuby2-expressing cells in ChAT-Cre mice injected with AAV-DIO-ChAT or AAV-DIO-mRuby2. Fluorescence intensity was normalized by non-transfected cells (mRuby2 negative). n=56 and 58 cells for mRuby2 and ChAT. n=3 animals/group. Welch’s t-test. **e,** Wheel running speed of mice that were injected with AAV-DIO-ChAT or AAV-DIO-mRuby2 at the cPPN. n=8 animals/group. Welch’s t-test. **f,g,** Rotarod (**f**) and balance beam (**g**, 4 mm rod) tests show that expression of mRuby2 did not affect enhancement of acquisition and maintenance of motor skills gained by running (mRuby2-Run vs. mRuby2-Ctrl), whereas exogenous ChAT expression blocked the enhancement (ChAT-Run vs. mRuby2-Run). The blockade was sustained after rest for 1 week. For immediate rotarod test, numbers of animals are 9 for mRuby2-Ctrl, 16 for ChAT-Ctrl, 9 for mRuby2-Run, 18 for ChAT-Run. For rotarod retest, numbers of animals are 9 for mRuby2-Ctrl, 6 for ChAT-Ctrl, 9 for mRuby2-Run, 7 for ChAT-Run. For immediate test of balance beam, numbers of animals are 10 for mRuby2-Ctrl, 9 for ChAT-Ctrl, 10 for mRuby2-Run, 9 for ChAT-Run. For balance beam retest, numbers of animals are 6 for mRuby2-Ctrl, ChAT-Ctrl, mRuby2-Run and 7 for ChAT-Run. ANOVA followed by Tukey’s test. NS, not significant; \**p*<0.05, \*\**p*<0.01, \*\*\**p*<0.001. Data shown are mean± SEM.

Mice that had run for 1 week, not subjected to behavioral tests, and allowed 1 week of rest now exhibited the same number of ChAT+ and GAD1+ neurons as control mice that had never run on a running wheel (Fig. 2k). No apoptosis or neurogenesis was detected in the cPPN of these mice (Supplementary Fig. 5c-f), indicating that the transmitter switch had spontaneously reversed. The time during which the transmitter switch persists (Fig. 2k) corresponds to the time during which the benefit of running on motor skill acquisition can occur (Fig. 1j,k), indicating a temporal correlation between transmitter switching and enhanced motor skill learning. This finding raised the possibility that the transmitter switch is necessary for running to enhance motor skill learning. Note that the transmitter switch reversed one week after running (Supplementary Fig. 7), even when training and testing on the rotorod and balance beam occurred immediately after running and the acquired motor skills persisted for at least 2 weeks (Fig. 1g-i). These results suggest that the transmitter switch is not necessary for maintaining the acquired motor skills.

**Fig. 7.**
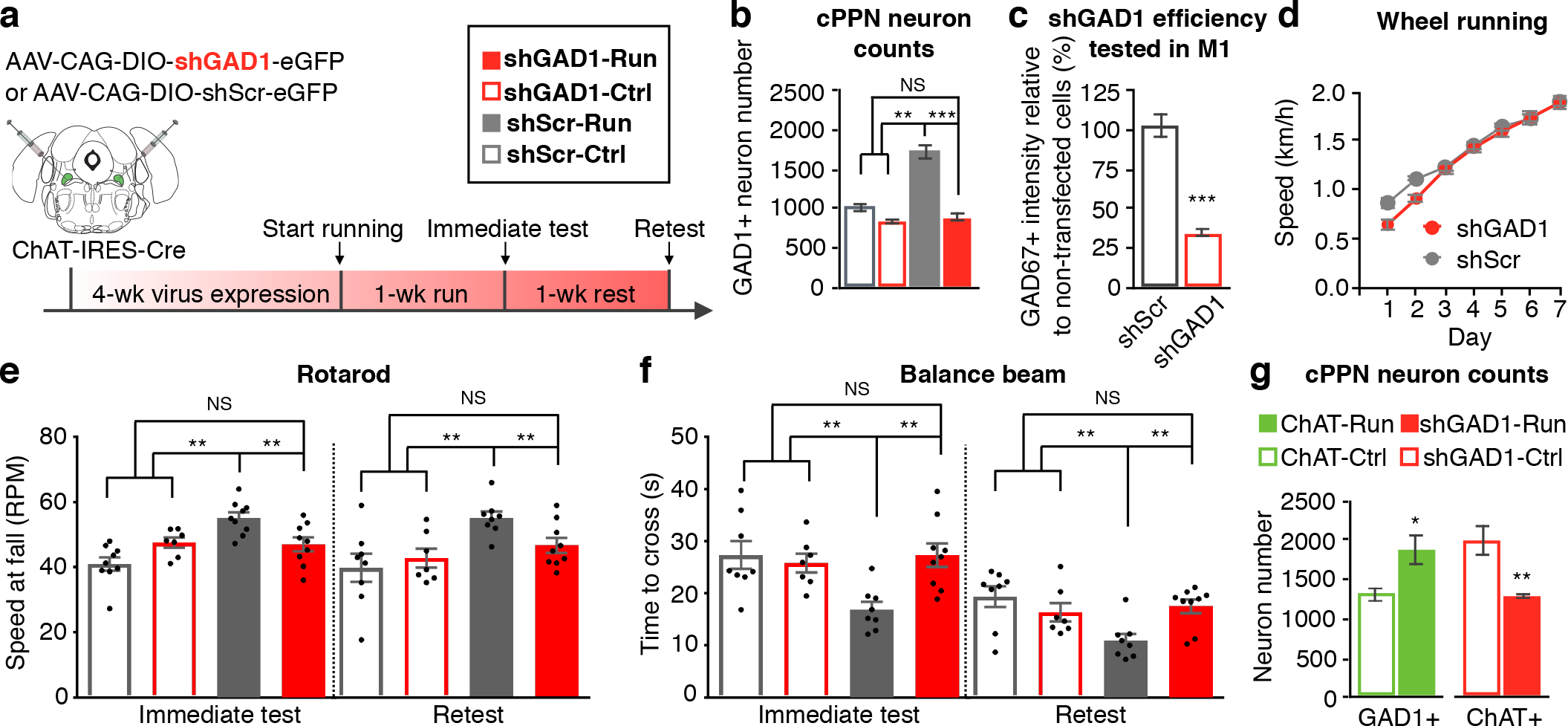
Gain of GAD1 in cholinergic cPPN neurons is also necessary for running-enhanced motor skill learning. **a,** Experimental design to override the gain of GAD1 in ChAT-Cre neurons and examine behavioral relevance. **b,** Numbers of GAD1 *in situ* stained neurons in the cPPNs of runners and non-running controls that were injected with AAV-DIO-shScr (scramble shRNA) or AAV-DIO-shGAD1 (shRNA for GAD1). n=7 for ChAT-Run, and 5 animals for the other three groups. Nonparametric Kruskal–Wallis test followed by Dunn’s correction. **c,** Fluorescence intensity of GAD67 immunoreactivity measured in M1 PV+ cells and data from PV+/GFP+ cells normalized by that from PV+/GFP-cells. n= 32 and 36 cells for shScr and shGAD1. n=3 animals/group. Welch’s t-test. **d,** Wheel running speed of ChAT-Cre mice that express AAV-DIO-shScr or AAV-DIO-shGAD1 in the cPPN. n=8 animals/group. Welch’s t-test. **e,f,** Rotarod (**e**) and balance beam (**f**, 4 mm rod) tests show that expression of the scramble shRNA did not affect the enhancement of acquisition and maintenance of motor skills gained by running (shScr-Ctrl vs. shScr-Run) whereas knocking down GAD1 blocked the enhancement (shGAD1-Run vs. shScr-Run). The blockade was sustained after rest for 1 week. Numbers of animals are 8 for shScr-Ctrl, 7 for shGAD1-Ctrl, 8 for shScr-Run, and 9 for shGAD1-Run. ANOVA followed by Tukey’s test. **g,** Numbers of GAD1 *in situ* stained neurons in the cPPN of 1-week runners and non-runner controls that express AAV-DIO-ChAT and numbers of ChAT immunostained neurons in 1-week runners and non-runner controls that express AAV-DIO-shGAD1. n=5 animals/group. Mann–Whitney U test. NS, not significant; \**p*<0.05, \*\**p*<0.01, \*\*\**p*<0.001. Data shown are mean± SEM.

To seek more direct evidence for neurotransmitter switching, we selectively tagged cholinergic neurons with genetic markers to reveal their change in transmitter identity following the exercise challenge. We used a well-characterized ChAT-Cre transgenic mouse line (*26,27*) that exhibited the same running-dependent loss of ChAT and gain of GAD1 expression as wild-type mice (Supplementary Fig. 8). We injected a Cre-dependent AAV vector (AAV-DIO-mRuby2) into the cPPN to permanently label cholinergic neurons with mRuby2 even when they have lost ChAT after running (Fig. 3a,b). We then scored the number of mRuby2+ neurons that express ChAT and/or GABA immunofluorescence and found that in control mice, 65% of cPPN cholinergic neurons tagged by mRuby2 expressed only ChAT, while 29% of them co-expressed ChAT and GABA, 4% expressed neither and 2% expressed only GABA (Fig. 3c). In runners, 24% of mRuby2+ neurons expressed only ChAT, 31% co-expressed ChAT and GABA, 12% expressed neither and 33% expressed only GABA (Fig. 3c). The decrease in neurons expressing only ChAT and increase in neurons expressing only GABA in the ChAT-Cre line identify the expression of GABA in formerly cholinergic neurons. More modest co-expression of ChAT and GAD has been observed in different transgenic mouse lines (*28*). The increase in number of neurons that express neither ChAT nor GABA suggests that switching neurons lose ChAT before gaining GABA.

**Fig. 8.**
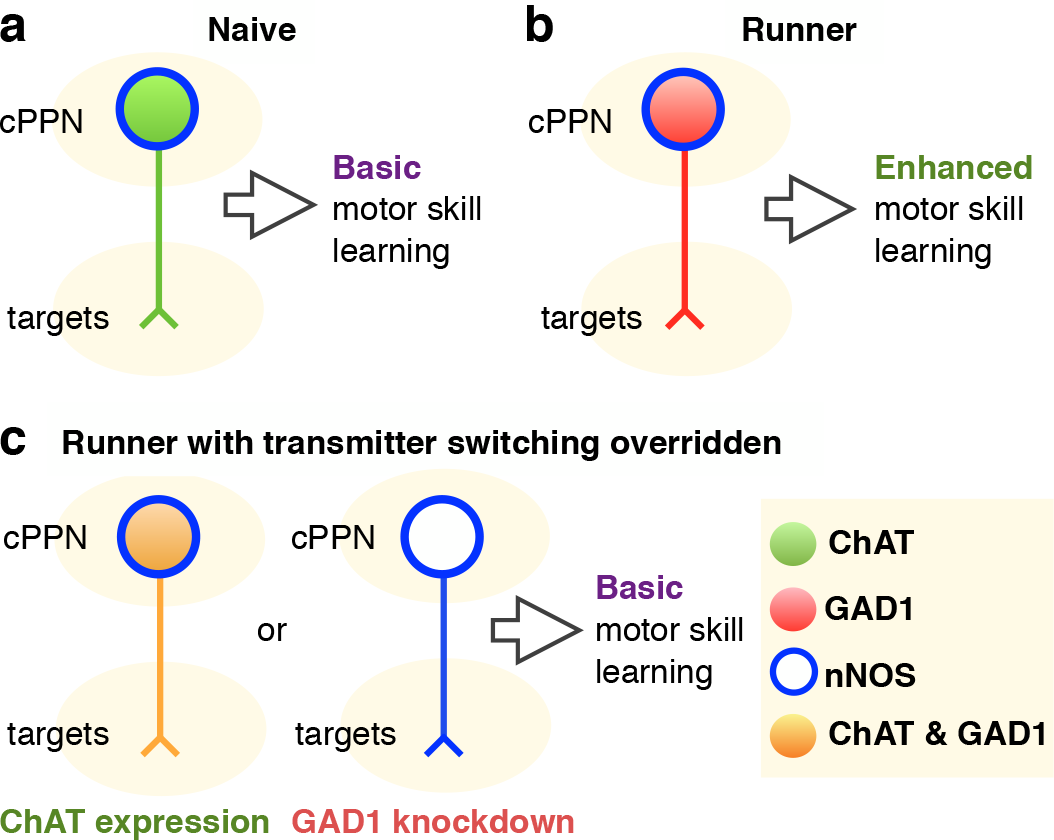
Transmitter switching in the cPPN regulates motor skill learning. **a,b,** Chronic running induces neurotransmitter switching from ACh to GABA in the cPPN and enhances motor skill learning. **c,** Both ChAT loss and GAD1 gain are necessary for running-enhanced motor skill learning.

### Switching involves regulation of transcript levels of transmitter synthetic enzymes

To understand whether the loss of ChAT and gain of GAD1 occur in the same population of neurons at the transcript level, we used sensitive fluorescence *in situ* hybridization to analyze the levels of mRNA of transmitter synthetic enzymes in cPPN neurons from runners and controls. We used the expression of neuronal nitric oxide synthase (nNOS) as a biomarker restricted to ChAT+ neurons in the PPN (*29*) (Fig. 4a,b). Although the number of ChAT+ neurons decreased with one week of running (Fig. 2g,h), the number of nNOS+ neurons did not change (Fig. 4c). We then combined immunofluorescent labeling of nNOS with RNAscope to detect mRNA encoding ChAT and GAD1 in nNOS+ neurons. By measuring the number of transcript puncta (Fig. 4d-g) and the total fluorescent area and fluorescence intensity per cell (Supplementary Fig. 9), we demonstrated a decrease in the number of ChAT transcripts and an increase in the number of GAD1 transcripts in neurons expressing nNOS in runners. While RNAscope revealed the presence of ChAT transcripts in all nNOS neurons in controls (Fig. 4e), immunostaining identified 11% of nNOS neurons that did not show ChAT immunoreactivity (Fig. 4b). These results imply that in nNOS neurons of controls, the bottom 11% of the distribution of ChAT mRNA puncta (Fig. 4e; expressing less than 8 puncta) lack detectable ChAT immunoreactivity. We next grouped neurons into the four categories of Figure 3 on the basis of ChAT and GAD1 mRNA expression (see Methods). In control mice, 40% of the cPPN nNOS+ cells were classic cholinergic neurons, while 39% of them co-expressed ChAT and GAD1, 6% expressed neither and 5% had switched from ChAT to GAD1 (Fig. 4e). However, in runners, the percentages changed to 18% for classic cholinergic neurons, 43% for co-expressing cells, 12% for neither and 27% that had switched (Fig. 4e). These changes in the four categories are comparable to those identified by immunocytochemistry (Fig. 3c), further supporting a switch from ChAT to GAD1 in nNOS neurons. The RNAscope assay reveals that switching involves up and down regulation of transcript levels and may not entail complete disappearance and de novo appearance of transcripts of transmitter synthetic enzymes.

### Switching neurons make projections to the SN, VTA and thalamus

To further test the link between neurotransmitter switching and motor skill learning, we identified targets innervated by the switching cholinergic neurons. PPN neurons project to the substantia nigra (SN), the ventral tegmental area (VTA) and the ventrolateral-ventromedial nuclei of the thalamus (VL-VM), all of which regulate motor skill learning (*30–35*). Anterograde tracing of mRuby2 in ChAT-Cre neurons (see Methods) followed by retrograde tracing with retrobeads demonstrated synaptic connections between cholinergic cPPN neurons and neurons in the SN, VTA and VL-VM (Fig. 5a-c and Supplementary Fig. 10). cPPN neurons originating projections to all three targets had lower mean numbers of ChAT transcript puncta in runners compared to controls, as demonstrated by triple labeling of retrograde beads, ChAT transcripts and nNOS protein (Fig. 5d-f and Supplementary Fig. 11). As noted above, nNOS neurons expressing less than 8 puncta appear to be those lacking ChAT immunoreactivity. In controls, the percentages of neurons that express less than 8 ChAT transcript puncta and project to the SN, VTA, and VL-VM were 11%, 8% and 6%, and increased to 42%, 11% and 19% in runners. Neurons in the increased percentages of cells expressing low numbers of ChAT-transcript puncta are likely to include those that switch transmitters. The greater increase for the SN suggests that cPPN neurons switching transmitters make a major projection to the SN with smaller projections to the VTA and VL-VM.

### Transmitter switching in the cPPN is necessary for runningFenhanced motor skill learning

To determine whether neurotransmitter switching is necessary for the beneficial effect of running on motor skill learning, we injected AAV-DIO-ChAT in the cPPN of ChAT-Cre mice to continuously express ChAT in all cholinergic neurons (Fig. 6a,b and Supplementary Fig. 12). The ChAT-Cre line (*26,27*) demonstrated the same running-dependent transmitter switch as wild-type mice, even with expression of control constructs (AAV-DIO-mRuby2, Fig. 6c; AAV-DIO-shScr, Fig. 7b). Overexpression of ChAT did not change the number of ChAT+ neurons in the cPPN of control ChAT-Cre mice and maintained the number of ChAT+ neurons at control levels in the cPPN after sustained running (Fig. 6c). Although the overexpression of ChAT caused a 3.3-fold increase in the level of ChAT expression in control mice (Fig. 6d and Supplementary Fig. 12c), it did not affect their basal motor skill learning (Fig. 6f,g and Supplementary Fig. 13e,f) or running activity in runner mice (Fig. 6e and Supplementary Fig. 13a,b). This may result from feedback inhibition of ChAT by ACh (*36,37*) that is likely to maintain ACh levels of the non-switching neurons in the physiological range. Mice that had received AAV-DIO-ChAT acquired the same running skill as wild-type mice or ChAT-Cre mice injected with AAV-DIO-mRuby2 but their motor learning on the rotarod and balance beam, tested directly after 1 week of running, was not enhanced (Fig. 6f,g). The slopes of the learning curves for both rotarod and balance beam behaviors were significantly steeper for runner mice expressing AAV-DIO-mRuby2 (10±2 rpm/trial and −3.9±1.0 s/trial) than for runners expressing the AAV-DIO-ChAT (6±1 rpm/trial, −1.3±0.4 s/trial; *p*=0.037 and *p*=0.035) and test performances of runners expressing AAV-DIO-mRuby2 were significantly better (Supplementary Fig. 13e,f). Overriding the loss of ChAT also prevented enhancement of motor skill learning when mice ran, were trained and tested, rested for one week and then re-tested (Fig. 6f,g). This finding makes it unlikely that exogenous expression of ChAT had simply delayed the improvement in motor skill learning.

Expression of a Cre-dependent viral construct (AAV-DIO-shGAD1) to suppress expression of GAD1 in the cPPN of ChAT-Cre mice similarly prevented improved motor learning following one week of running, compared to mice expressing a Cre-dependent scrambled shRNA sequence (AAV-DIO-shScr; Fig. 7a-f, Supplementary Figs. 13 and 14). Knockdown of GAD1 maintained the number of GAD1+ neurons at control levels in the cPPN after sustained running (Fig. 7b). Because immunostaining of GAD67 in the PPN detects a large number of synaptic puncta that makes it challenging to analyze cell bodies, the knockdown efficiency of GAD67 expression by GAD1 shRNA was tested in the motor cortex and measured to be 65% (Fig. 7c and Supplementary Fig. 14a,b). Suppressing the gain of GAD1 did not affect acquisition of the wheel running skill (Fig. 7d and Supplementary Fig. 13c,d) but motor learning on the rotarod and balance beam, tested directly after one week of running, was not enhanced for a period that was extended to one week of rest (Fig. 7e,f). The slopes of the learning curves were again steeper for runner mice expressing AAV-DIO-shScr (9±1 rpm/trial, −6.7±0.9 s/trial) than for runners expressing AAV-DIO-shGAD1 (5±2 rpm/trial, −3.7±1.0 s/trial; p = 0.073 and p=0.041) and test performances for runners expressing AAV-DIO-shScr were again significantly better (Supplementary Fig. 13g,h). These results suggest that both the loss of ACh and gain of GABA are required for enhancement of motor skill learning. Overriding the loss of ChAT or suppressing the gain of GAD1 did not affect running-induced activity in cholinergic cPPN neurons (Supplementary Fig. 15). Notably, suppressing the gain in GAD1 expression did not affect the loss of ChAT expression and vice versa (Fig. 7g). Although expression of these two transmitter synthetic enzymes is inversely correlated, the two are not reciprocally regulated.

## Discussion

Our findings provide new insight into the mechanism by which sustained running improves acquisition of motor skills (Fig. 8). The functional significance of ACh-to-GABA transmitter switching is demonstrated by changes in behavior that are reversed by overriding the switch. Switching involves changes in levels of transcripts of transmitter synthetic enzymes. Activity-dependent transcription factor phosphorylation (*13,14*) and microRNA regulation (*38*), which have been implicated in transmitter switching in the developing nervous system, are candidates for implementing the switch in the adult CNS. Transmitter switching, particularly when it can change the sign of the synapse from excitatory to inhibitory, appears to rewire motor circuitry to enhance motor skills. The persistence of learned behaviors after the transmitter switch has reversed implies that there is continued capacity for plasticity in locomotor circuitry.

We examined three targets of the cPPN and found that cholinergic cPPN neurons projecting to each of the three targets showed a significant reduction of ChAT transcripts (Fig. 5). This suggests that the reduction of ChAT expression might be observed in cholinergic cPPN neurons projecting to other targets as well. Multi-target regulation is not surprising because cholinergic PPN neurons have an average of five axonal collaterals that allow a single cholinergic neuron to project to many targets (*18*). Cholinergic cPPN neurons may regulate motor function and other behaviors by gating the activity of downstream targets through transmitter switching.

Although dopaminergic and noradrenergic neurons are implicated in motor function plasticity (*39–42*), our results unexpectedly indicate that acquisition of high-demand rotarod and balance beam performance requires transmitter switching in cholinergic cPPN neurons. Overriding transmitter switching did not affect the ability to run on a running wheel (Figs. 6 and 7), perhaps because running is not an exceptionally taxing motor skill. Consistent with these results, selective lesions of cholinergic PPN neurons impair learning of high-demand running on an accelerating-speed rotarod but do not affect learning of low-demand running on a fixed-speed rotarod or basal locomotion (*43*). In contrast, glutamatergic and GABAergic PPN neurons regulate gait and speed of locomotion (*44–46*).

The balance of Parkinson’s disease and stroke patients is improved following sustained treadmill training (*47,48*). Our finding that transmitter switching after running is a critical event for improving motor skill learning suggests that transmitter switching may be important in many circumstances where sustained exercise benefits behavior.

## Acknowledgements

We thank Kyle Jackson, Vaidehi Gupta and Wuji Jiang for experimental assistance, Alex Glavis-Bloom for technical support and Larry Squire and Takaki Komiyama for comments on the manuscript. We thank all members of the Spitzer lab for suggestions on experimental design and the manuscript. We thank Stefan Leutgeb and Byungkook Lim for providing transgenic mice. We thank Jennifer Santini and the UCSD Microscopy Core for technical support with imaging and Amanda Roberts and the Scripps Animal Models Core for suggestions on behavior experiments. This research was supported by grants to N.C.S from the Ellison Medical Foundation, the W. M. Keck Foundation and the Overland Foundation.

## Author Contributions

H.L. and N.C.S. conceived the study, designed the experiments, interpreted the results and wrote the paper. H.L. performed the experiments and analyses.

## Competing interesting statement

The authors declare that they have no competing financial interests.

## Methods

### Mice

All animal procedures were carried out in accordance with NIH guidelines and approved by the University of California, San Diego Institutional Animal Care and Use Committee or Scripps Institutional Animal Care and Use Committee. C57BL/6J (JAX#000664) mice were obtained from Jackson Laboratories. ChAT-IRES-Cre (JAX#006410) mice were obtained from the Byungkook Lim lab and Jackson Laboratories. PV-IRES-Cre (JAX#008069) mice were provided by the Stefan Leutgeb lab. Animals were maintained on a 12 h:12 h light:dark cycle (light on: 10:00 pm-10:00 am) with food and water ad libitum. The ChAT-IRES-Cre colony was maintained by breeding homozygous male ChAT-Cre mice with female wild-type C57BL/6J mice. Heterozygous ChAT-Cre offspring were used in the study. Both heterozygous and homozygous PV-IRES-Cre mice were used. All experiments were performed on 8 to 12-week-old male mice.

### Behavioral assays

#### Wheel running

Mice were single-housed in hamster cages and provided with FastTrac (Bio-serv, K3250) or digital (Med Associates ENV-044) running wheels that are identical in shape and size. Mice were allowed voluntary running for one week and were continuously recorded with Swann DVR4-2600 infrared video cameras. Control mice were housed with running wheelbases without wheels. Episode duration and time with the wheel were hand-scored using JWatcher software. The maximum angular excursion of mouse movements on the running wheels was plotted using Image J and measured with a protractor. The digital running wheels recorded running distance and running speed was calculated as distance divided by time.

#### Rotarod

Training and tests were performed as the rotarod (Ugo Basile, 57624) accelerated from 5 rpm to 80 rpm in 6 min. The rpm at which mice fell off was recorded by the rotarod. Mice were trained for 9 trials on the first day, followed by 3 tests the next day and 3 retests after 1, 2, or 4-weeks rest. There was a 10-minute interval between each trial/test. The single digit accuracy of the slopes of learning curves reflects the accuracy of recording the rpm at fall by the rotarod.

#### Balance beam

Mice were trained and tested with balance beams 1 meter long and 0.75 meters above the floor. 12 mm and 6 mm square beams (S-12 and S-6) were single plastic horizontal bars while 6 mm and 4 mm rod beams (R-6 and R-4) consisted of two stainless steel parallel bars 5 cm apart. On the training day mice were trained to cross S-12 for 3 trials, S-6 for 3 trials, R-6 for 3 trials and then R-4 for 3 trials, each 10 minutes apart. On the test and retest day mice were allowed to cross all four beams in the same order, three times for each, 10 minutes apart. A video camera was installed at the end of the beam where mice started walking. At the other end of the beam, a black box with an entry facing the beam was installed to attract mice to cross the beams. Time to cross the beam and enter the escape box was hand-scored and the investigator was double-blinded to the history of the mice. The slopes of learning curves are accurate to one decimal place because the time to cross the beam was hand scored and human response time is 0.1~0.2 seconds.

#### Motor skill learning

The mean speed at fall from a rotarod for the nine trials on the training day was fitted by a one-phase association model. Mean data points for each training trial were plotted and fitted using GraphPad Prism 7 software and the coefficient of determination (R^2^) was used to justify the fit (R^2^>0.96 for controls and R^2^>0.95 for runners, with MATLAB). The mean time to cross a 4 mm rod beam for the three trials on the training day was fitted by linear regression (R^2^>0.98 for controls and R^2^>0.99 for runners, with MATLAB).

#### Locomotor activity

Mice were tested for 120 minutes in polycarbonate cages (42 × 22 × 20 cm) placed in frames (25.5 × 47 cm) mounted with two levels of photocell beams at 2 and 7 cm above the bottom of the cage (San Diego Instruments, San Diego, CA). The two sets of beams detect both horizontal (roaming) and vertical (rearing) behavior. A thin layer of bedding material covered the bottom of the cage. Data were collected in 1-min epochs.

### Histology and immunocytochemistry

Animals were perfused transcardially with phosphate-buffered saline (PBS) followed by 4% paraformaldehyde (PFA) in PBS right after the last episode of running, i.e. mice started to run at 10 am at light-off and kept running until they were perfused between 2 to 3 pm. Brains were dissected and post-fixed in 4% PFA for 16 to 24 h at 4°C, washed in PBS for 1 min and transferred to 30% sucrose in PBS for 2 days at 4°C. 40-μm coronal sections were cut on a microtome (Leica SM2010R) and stained.

For immunostaining, sections were permeabilized and blocked in 24-well culture plates for 2 h in a blocking solution (5% normal horse serum, 0.3% Triton X-100 in PBS) at 22-24°C. Primary and secondary antibodies were diluted in the blocking solution. Incubation with primary antibodies was performed for 48 h on a rotator at 4°C. After washing in PBS (3 times, 15 min each), secondary antibodies were added for 2 h at 22-24°C. For immunofluorescence, sections were mounted with Fluoromount-G (Southern Biotech) or ProLong Gold Antifade Mountant (Life Technologies) containing DRAQ-5 (Thermo Fisher, 62251, 1:1000 dilution; when nuclear staining was needed) after washes in PBS (3 times, 15 min each).

For DAB (3,3’-Diaminobenzidine) staining, sections were treated with 0.3% hydrogen peroxide for 30 min, washed in PBS (3 times, 5 min each), incubated with Vectastain Elite ABC HRP mixture (Vector Laboratories, PK-6100) for 45 min, washed in PBS (3 times, 15 min each), and signals were developed using the DAB Peroxidase Substrate kit (Vector Laboratories, SK-4100).

Primary antibodies used in this study were goat anti-ChAT (Millipore, AB144P, 1:500), rabbit-anti-nNOS (Thermo Fisher, 61-7000, 1:500), goat anti-cFos (Santa Cruz, sc-52G, 1:300), rabbit-anti-cFos (Santa Cruz, sc-52, 1:300), mouse-anti-cFos (Abcam, ab208942, 1:500), rabbit-anti-PV (Swant, PV25, 1:2000), goat-anti-VAChT (Millipore, ABN100, 1:500), mouse-anti-NeuN (Millipore, MAB377, 1:500), rabbit-anti-GABA (Sigma-Aldrich, A2052, 1:1000) rabbit-anti-GFP (Thermo Fisher, A11122, 1:1000), chicken anti-GFP (Abcam, ab13970, 1:1000), guinea pig anti-GFP (Synaptic Systems, 132005, 1:3000), goat-anti-doublecortin (Santa Cruz, sc-8066, 1:300) and rabbit-anti-Ki67 (Cell Signaling, 9129, 1:300). Secondary antibodies for immunofluorescence were from Jackson ImmunoResearch Labs and used at a concentration of 1:600: Alexa Fluor-488 donkey-anti-rabbit (705-545-003), Alexa Fluor-488 donkey-anti-guinea pig (706-545-148), Alexa Fluor-488 donkey-anti-mouse (715-545-150), Alexa Fluor-488 donkey-anti-goat (705-545-147), Alexa Fluor-594 donkey-anti-goat (705-585-147), Alexa Fluor-594 donkey-anti-mouse (715-585-150), Alexa Fluor-647 donkey-anti-goat (705-605-147) and Alexa Fluor-647 donkey-anti-rabbit (711-605-152). Biotinylated goat anti-rabbit (BA-1000) and horse anti-goat (BA-9500) secondary antibodies for DAB staining were from Vector Laboratories and used at a concentration of 1:300.

### *In situ* hybridization

PFA fixed brains were dissected, post-fixed in 4% PFA for 16 to 24 h at 4°C and transferred to 30% DEPC-sucrose in PBS for 2 days. Subsequently, brains were embedded in 30% DEPC-sucrose and frozen with dry ice. 40-μm cryosections were collected on Superfrost Plus slides (VWR, 48311-703) and used for mRNA *in situ* hybridization. Complementary DNAs (cDNAs) of *gad1* or *slc17a6* (sequence from Allen Brain Atlas) were cloned in ~800-base-pair segments into a pGEM vector. Antisense complementary RNA (cRNA) probes were synthesized with T7 (Promega, P2075) or Sp6 polymerases (Promega, P1085) and labeled with digoxigenin (Roche, 11175025910). Hybridization was performed with 1 to 5 μg/ml cRNA probes at 65 °C for 20 to 24 h. Probes were detected using Anti-Digoxigenin-AP Fab fragments (Roche, 11093274910, 1:5000). Signals were developed using a mixture of 4-Nitro blue tetrazolium chloride (Roche, 11383213001) and BCIP 4-toluidine salt solution (Roche, 11383221001).

Fluorescent RNAscope *in situ* hybridization was performed according to the manufacturer’s instructions (Advanced Cell Diagnostics) with some modifications: In an RNase-free environment, 12-μm fixed brain sections were mounted on Superfrost Plus slides immediately after microtome sectioning and air-dried in a 60°C oven for 30 min. Sections were rehydrated in PBS for 2 min and incubated for 5 min in 1× target retrieval solution at 95°C. Sections were then rinsed with distilled water for 5 seconds and rinsed in 100% ethanol for 5 seconds. After air-drying, sections were incubated with the following solutions in a HybEZ humidified oven at 40°C with three rinsing steps in between each: protease III, 30 min; probes, 2 h; amplification (Amp) 1-fluorescence (FL), 30 min; Amp 2-FL, 15 min; Amp 3-FL, 30 min; and Amp 4-FL, 15 min. Ready-to use Amp 1-FL, Amp 2-FL, Amp 3-FL, Amp 4-FL and 50x washing solution for rinsing steps were included in the RNAscope Multiplex Fluorescent Reagent kit. Standard immunofluorescent staining was subsequently performed in the dark as previously described (*15*). Probes for mouse *chat* mRNA and *gad1* mRNA were from Advanced Cell Diagnostics. Sections were 6 to 7 μm thick post-processing. Six optical sections of each physical section were examined and regions of interest (ROIs) were drawn around the boundaries of nNOS cells on the optical section that showed the best focal plane for each cell (largest cross-sectional area). These ROIs were scored for ChAT and GAD1 transcript puncta using Image J. The average area of ROIs was consistent between the control and runner groups. Based on the expression of ChAT and GAD1 transcript puncta, the neurons were grouped into four categories 1) classic cholinergic neurons: ⩾ 8 ChAT transcript puncta, no GAD1 transcript puncta; 2) co-expressing neurons: ⩾8 ChAT puncta, ⩾1 GAD1 puncta); 3) neurons expressing neither: <8 ChAT puncta, no GAD1 puncta; 4) switched neurons: <8 ChAT puncta, ⩾1 GAD1 puncta.

### Birthdating

Mice were intraperitoneally injected with BrdU (50 mg/kg) once every 12 hr for 1 week. PFA-fixed brains were dissected, post-fixed and dehydrated as described above. 40-μm cryosections were collected and treated with 1 M HCl for 30 min at 45°C for DNA denaturation. After rinsing in PBS (3 times, 5 min each), sections were incubated in a mixture of 0.3% Triton X-100 and 5% horse serum in PBS for 1 h, incubated in rat-anti BrdU antibody (AbD Serotec, MCA2060, 1:300) at 4°C overnight, rinsed in PBS (3 times, 5 min each) and amplified by Alexa Fluor-488 donkey anti-rat antibody (Life Technologies, A21208, 1:600). Sections were mounted with Fluoromount containing DRAQ-5 (1:1000).

### TUNEL assay

The *In Situ* Cell Death Detection (TUNEL) Kit with TMR Red (Roche, 12156792910) was used to detect *in situ* apoptosis. 40-μm cryosections were re-fixed with 1% PFA for 20 min at 22-24°C and rinsed with PBS (3 times, 5 min each). Sections were then permeabilized in 0.1% sodium citrate and 1% Triton X-100 for 1 h at 22-24°C. After rinsing in PBS (3 times, 5 min each), sections were incubated with TUNEL reaction solution according to the vendor’s instruction, i.e. incubated in a mixture of 25 μL of terminal-deoxynucleotidyl transferase solution and 225 μL of label solution. Incubation was performed in a humidified chamber for 3 h at 37°C in the dark. Sections were rinsed and mounted with Fluoromount containing DRAQ-5 (1:1000). For a positive control, sections were treated with DNase I (10 U/mL, New England Biolabs, M0303S) for 1 h at 37°C and rinsed in PBS (3 times, 5 min each), followed by incubation with the TUNEL mixture.

### Imaging and data analysis

Fluorescent images were acquired with a Leica SP5 confocal microscope with a 25x/0.95 water-immersion objective and a z resolution of 1 μm. Leica Application Suite X software was used for fluorescent cell counting and all sections within the confocal stacks were examined without maximal projection. RNAscope *in situ* signals were quantified using Image J. Exemplar images are maximum intensity projections of 6 consecutive confocal sections when RNAscope signals are included (Figs. 4d and 5e) and 16 to 24 confocal sections for other immunofluorescence images. Exemplar images for DAB staining and *in situ* hybridization were acquired with a NanoZoomer Slide scanner (Hamamatsu S360) with a 20x/0.75 air objective. Counting DAB-stained c-fos neurons per unit area was performed with Image J on images harvested from the NanoZoomer. To delimit the PPN in sections stained for c-fos (Fig. S2), or for TUNEL and Ki67/DCX (apoptosis and neurogenesis, Fig. S5), an adjacent section was stained for ChAT to define the boundaries of the PPN.

### Stereological counting

Stereo Investigator software (MBF Bioscience) was used to count DAB immunostained or *in situ* hybridization-stained cells. Counting was carried out using optical fractionator sampling on a Zeiss Axioskop 2 microscope (40x/0.65 Ph2 objective) equipped with a motorized stage. The population of midbrain PPN neurons was outlined on the basis of ChAT immunolabeling, with reference to a coronal atlas of the mouse brain (Franklin and Paxinos 2007, 3^rd^ edition) and anatomical landmarks such as fiber tracts. “Caudal PPN” refers to the caudal half of PPN with reference to a transverse line normal to and halfway along the rostrocaudal axis; “rostral PPN” refers to the rostral half. To count GABAergic and glutamatergic neurons in the sections stained by *in situ* hybridization, an adjacent section was stained for ChAT to define the ROI for the boundaries of PPN. The area of the ROI and the cell density were compared between the control and experimental groups to ensure that the counted areas were comparable between the two groups and that changes in cell density were consistent with changes in the absolute counts. Pilot experiments with continuous counting determined that counting every other section was sufficient to estimate the number of cholinergic, GABAergic, glutamatergic and nitroxidergic neurons in the cPPN of each brain. Consequently, five to six sections were counted for each mouse brain. The average section thickness was measured prior to counting. Sections shrank from 40 μm to 22–24 μm after staining and dehydration. Exhaustive counting was performed and no sampling grid was skipped because the distribution of neurons in the cPPN is uneven. Counting was performed by two investigators double-blinded to the origin of the sections.

### Viral constructs and injection

Seven to eight-week-old male mice were anesthetized with a mixture of 120 mg/kg ketamine and 16 mg/kg xylazine and head-fixed on a stereotaxic apparatus (David Kopf Instruments Model 1900) for all stereotaxic surgeries. The caudal PPN was targeted bilaterally (cPPN: anterior–posterior (AP), −4.80 mm from bregma; mediolateral (ML), ±1.25 mm; dorsal–ventral (DV), −3.25 mm from the dura). A total of 300 nL of the Cre-on viral vectors were injected into each cPPN of ChAT-Cre mice at a rate of 100 nL/min using a syringe pump (PHD Ultra™, Harvard apparatus, no. 70-3007) installed with a microliter syringe (Hampton, no.1482452A) and capillary glass pipettes with filament (Warner Instruments, no. G150TF-4). Pipettes were left in place for 5 min after injection. Titers of recombinant AAV vectors ranged from 7.1 × 10^12^ to 2.2 × 10^13^ viral particles/ml, based on quantitative PCR analysis. pAAV-hSyn-DIO-ChAT-P2A-mRuby2 and pAAV-hSyn-DIO-mRuby2 plasmids were constructed in our lab and AAV2/8 vectors were packaged in the Salk Viral Vector Core. Vector Biolabs produced AAV8-CAG-DIO-shRNAmir-mGAD1-eGFP and AAV8-CAG-DIO-shRNAmir-Scramble-eGFP vectors. The shRNA sequence for mouse GAD1 is *5’+GTCTACAGTCAACCAGGATCTGGTTTTGGCCACTGACTGACCAGATCCTTTGACTGTAGA+3’.*

### Tract tracing

For anterograde tracing, 300 nL of Cre-on AAV8-DIO-mRuby2 vectors were unilaterally injected into the cPPN of ChAT-Cre mice as described above and axonal tracts were imaged by Leica confocal microscope. For retrograde tracing, a volume of 100 nL of retrobeads (Red Retrobeads™ IX or Green Retrobeads™ IX, Lumafluor) was injected at 100 nL/min into the SNc or VTA and VL-VM. Pipettes were left in place for 10 min after injection. 3 weeks after injection, mice were sacrificed for brain examination or housed with running wheels to examine running-induced loss of ChAT transcripts using RNAscope combined with nNOS immunostaining. ChAT transcripts in neurons that contained retrobeads were quantified using Image J. Retrobead infusion coordinates were SNc (from bregma AP −3.20 mm; ML ±1.50 mm; and DV from the dura, −4.00 mm), VTA (AP, −3.10 mm; ML, ±0.50 mm; and DV, −4.25 mm), and VL-VM (AP, −2.20 mm; ML, ±1.75 mm; and DV, −3.25 mm).

### Statistics and reproducibility

For comparisons between two groups, data were analyzed by Welch’s t-test for normally distributed data and the Mann–Whitney U test for data not normally distributed, using Graph Pad Prism 7. For repeated samples, data were analyzed by paired t-test. Statistical analyses of the data were performed using Prism 7 software for the number of animals for each experiment indicated in the figure legends. Means and SEMs are reported for all experiments. For comparisons between multiple groups, ANOVA followed by Tukey’s test was used for normally distributed data. The nonparametric Kruskal–Wallis test followed by Dunn’s correction was used for data that were not normally distributed. Experiments were carried out independently twice (Figs. 2b,c, 4a,b, 5b,c, 7b,c and Supplementary Figs. 1f and 5), three times (Figs. 1b,c, h-j, 2d-k, 3, 4c, d-h, 5e,f, 6c,d, 7g, and Supplementary Figs. 1a-d, 2, 3, 4, 6, 7, 8, 9, 11b-d, 12, 13a-d, 14c, 15), or four or more times (Figs. 1c-f, 6e-g, 7d-f and Supplementary Figs. 13e-h).

