## Supplementary Figures for "Exercise enhances motor skill learning by neurotransmitter switching in the adult midbrain"

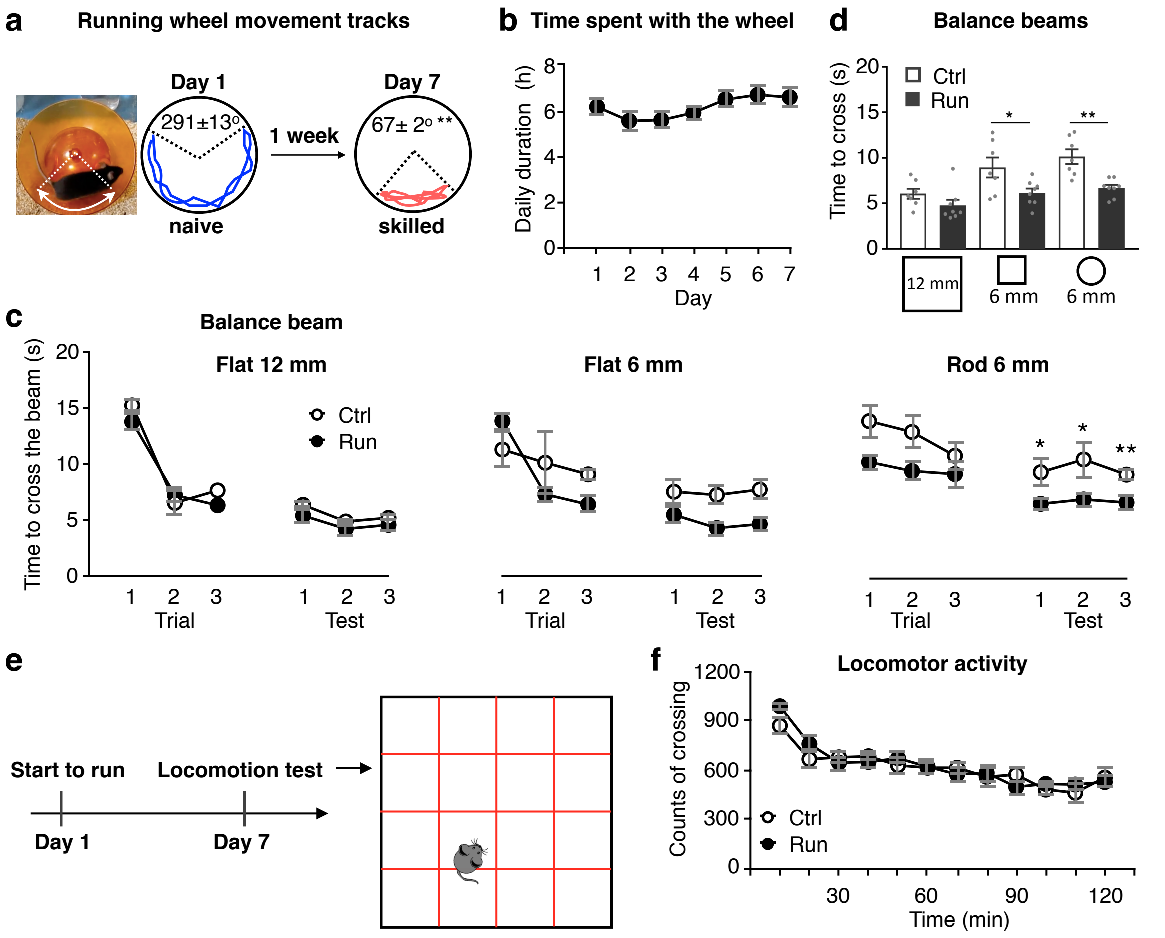


**Supplementary Figure 1**

**Running enhances motor skill learning but does not affect basic locomotor activity.**

**a,** Image of a runner mouse and examples of mouse movement tracks on the running wheel. Angles are mean mouse movements subtended while running. n=6 tracks from 3 animals/group. **b**, Daily duration that mice spent with running wheels during one week of running. n=8 animals/group. **c,** Time to cross a 1 meter long, 0.75 meter high balance beam during each trial of training or each test on the day after training. Beam shape (flat or rod) and size (diameter) are indicated. **d,** Mean time to cross the balance beams in three tests on the day after training. For (**c**,**d**), n=7 animals for Ctrl and 8 animals for Run. **e,** Experimental design for locomotion test. **f**, Average counts of mice crossing laser beams in the novel locomotion chamber (**e**) in a 2-hour test. n=6 animals/group. Statistical significance *$p$<0.05, **$p$<0.01 was assessed by Welch's t-test. NS, not significant. Data shown are mean± SEM.

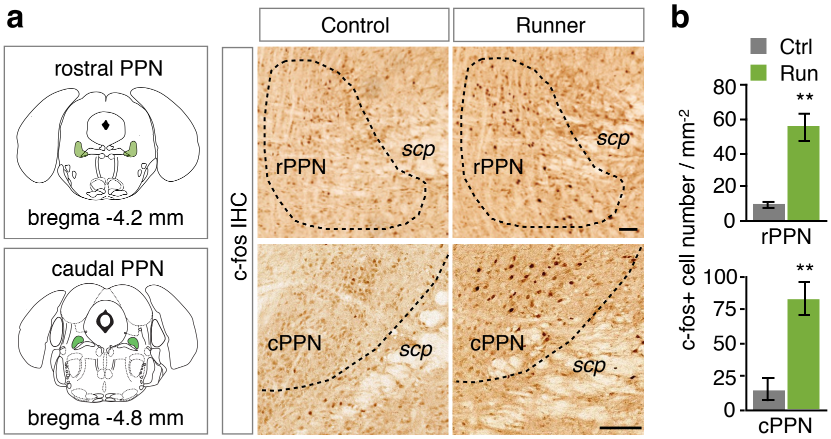


**Supplementary Figure 2**

**Running activates both rostral and caudal PPN.**

**a,** Left panels show the PPN (green) in coronal brain sections. Middle and right panels illustrate 3,3'-diaminobenzidine (DAB) immunohistochemistry of c-fos in rostral PPNs (rPPN, upper) or caudal PPNs (cPPN, lower) of controls and 1-week runners. Dotted lines outline the PPN. Dark brown stain indicates c-fos+ neurons. Scale bar, 100 µm. **b,** Counts of c-fos+ neurons in the rostral and caudal PPN. n=6 animals/group. Statistical significance ***p*<0.01 was assessed by Mann–Whitney U test. Data shown are mean± SEM.

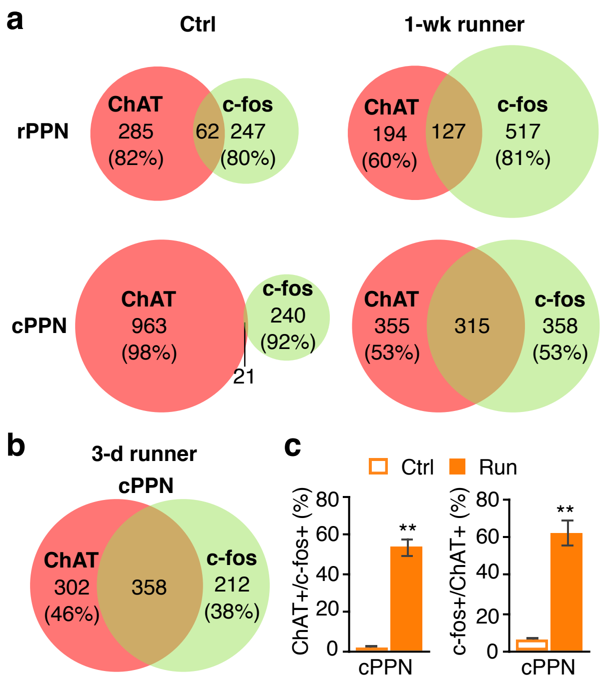


**Supplementary Figure 3**

**A substantial fraction of PPN c-fos+ neurons are cholinergic and cPPN cholinergic activation is more relevant to running.**

**a**, Co-localization of ChAT and c-fos in the PPN of controls and 1-wk runners. Cell numbers and percentages of cholinergic (red) and non-cholinergic (green) neurons are shown. n=3 animals/ group. **b**, Co-localization of ChAT and c-fos in the caudal PPN of mice that had run for 3 days. Cell numbers and percentages are shown. n=3 animals. **c**, The percentage of the ChAT+, c-fos+ neurons in the ChAT+ neuron population (left) or in the c-fos+ neuron population (right) in the cPPN of 3-day runners and controls. n=3 animals/group. Statistical significance $^{**}p<0.01$ was assessed by Mann–Whitney U test. Data shown are mean± SEM.

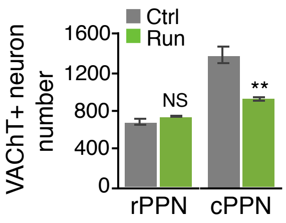


**Supplementary Figure 4**

**Running reduces the number of VAChT+ neurons in the cPPN but not in the rPPN**.

Stereological counts of VAChT+ neurons. n=6 animals/group. Statistical significance ***p*<0.01 was assessed by Mann–Whitney U test. NS, not significant. Data shown are mean± SEM.

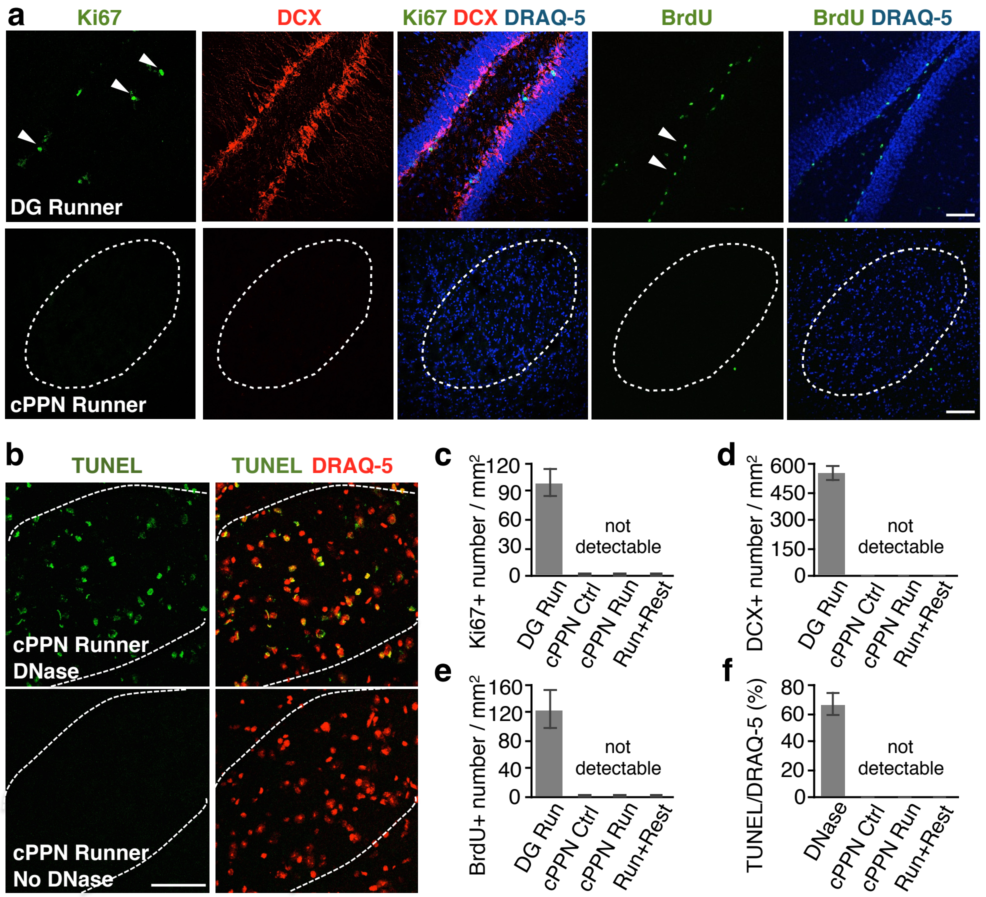


**Supplementary Figure 5**

**Apoptosis and neurogenesis are not apparent in the caudal PPN of runner mice.**

**a,** Sections of dentate gyrus (DG, upper panels) and cPPN (lower panels) from 1-week runners triple-stained for Ki67, DCX, and DRAQ-5 (nuclear marker) (Columns 1-3) or double-stained for BrdU and DRAQ-5 (columns 4-5). For BrdU labeling, mice were i.p. injected with BrdU (50 mg/kg) once every 12 hr for 1 week. Dotted lines outline the cPPN. Arrowheads point to Ki67+ (upper panel in column 1) or BrdU+ cells (upper panel in column 4). Scale bar, 100 µm. **b,** Sections of the cPPN of a 1-wk runner are double-stained for TUNEL and DRAQ-5. Dotted lines outline the cPPN. Upper panels: DNase-treated tissue as positive control. Lower panels: no DNase treatment. Scale bar, 100 µm. **c**-**e,** Statistical summary of (**a**) combined with littermate non-runner controls and mice that ran for 1 week followed by 1 week of rest. DG Run, dentate gyrus of a runner mouse. The region of interest that was quantified includes only the granule layer and not the hilus of the dentate gyrus. n=3 animals, 12 sections per group. **f,** Statistical summary of (**b**) combined with littermate non-runner controls and mice that ran 1-week followed by 1 week of rest. n=3 animals, 12 sections per group. Data shown are mean± SEM.

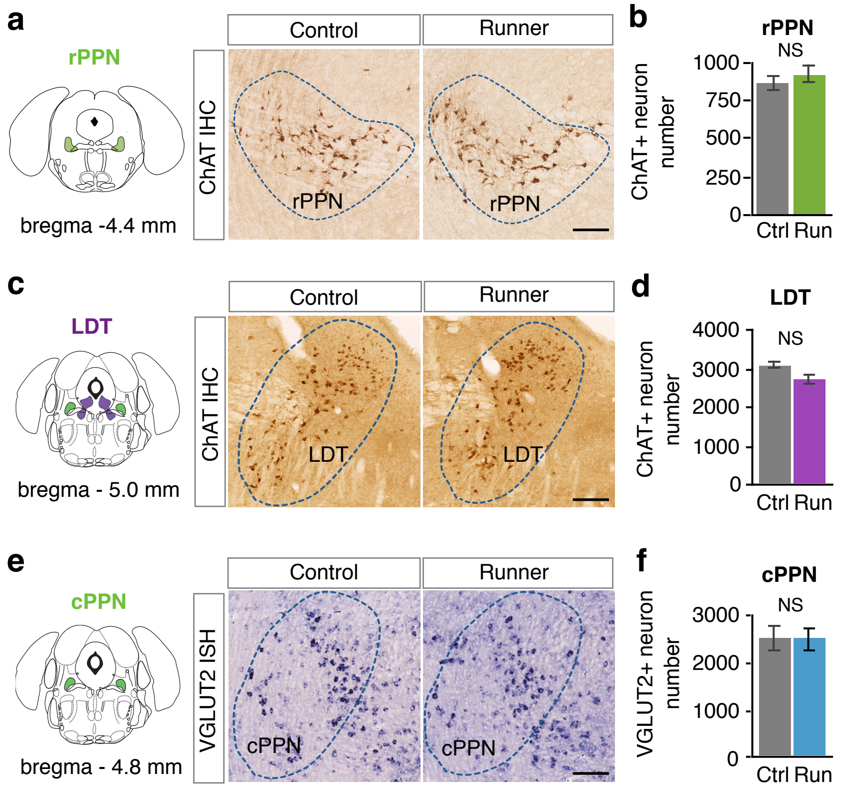


**Supplementary Figure 6**

**Running does not change the number of ChAT+ neurons in the rostral PPN (rPPN) or the laterodorsal tegmental nucleus (LDT) or the number of VGLUT2+ neurons in the caudal PPN (cPPN).**

**a,** The left panel shows the rPPN (green) in a coronal brain section. Middle and right panels illustrate DAB staining of ChAT in rPPN of a control and 1-week runner. Dotted lines outline the rPPN. Dark brown stain indicates ChAT+ neurons. Scale bar, 100 µm. **b,** Stereological counts of ChAT+ neurons in the rostral PPN. n=6 animals/group. **c,** The left panel shows the LDT (purple) and the caudal PPN (green) in coronal brain sections. Middle and right panels illustrate DAB staining of ChAT in LDT of a control and 1-week runner. Dotted lines outline the LDT. Dark brown stain indicates ChAT+ neurons. Scale bar, 100 µm. **d,** Stereological counts of ChAT+ neurons in the LDT. n=4 animals/group. Scale bar, 100 µm. **e,** The left panel shows the cPPN (green) in a coronal brain section. Middle and right panels illustrate *in situ* hybridization staining of VGLUT2 of a control and a 1-week runner. Dotted lines outline the cPPN. Dark blue-purple stain indicates VGLUT2+ neurons. Scale bar, 100 µm. **f,** Stereological counts of VGLUT2+ neurons in the cPPN. n= 4 animals for Ctrl and 5 animals for Run. Statistical significance was assessed by Mann–Whitney U test. NS, not significant. Data shown are mean± SEM.

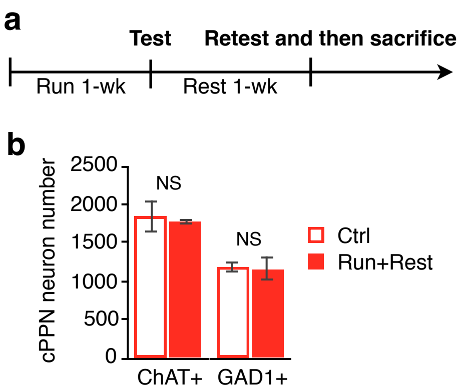


**Supplementary Figure 7**

**Running-induced transmitter switching reverses after 1 week of rest, even though mice experienced behavioral tests immediately after running and retests one week later.**

**a,** Experimental design. **b,** cPPNs from Ctrl or “Run+Rest” mice of the “1-wk rest group” (**Fig. 1g,h**) were stained for ChAT or for GAD1. n=5 animals/group. Statistical significance was assessed by Mann–Whitney U test. NS, not significant. Data shown are mean± SEM.

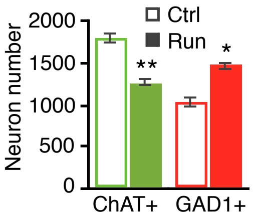


**Supplementary Figure 8**

**Running for 1 week decreases the number of ChAT+ neurons and increases the number of GAD1+ neurons in the caudal PPN of a ChAT-IRES-Cre mouse line.**

ChAT was detected by DAB staining. GAD1 was detected by *in situ* hybridization. n=5 animals/group. Statistical significance *$p$<0.05, **$p$<0.01 was assessed by Mann–Whitney U test. NS, not significant. Data shown are mean± SEM.

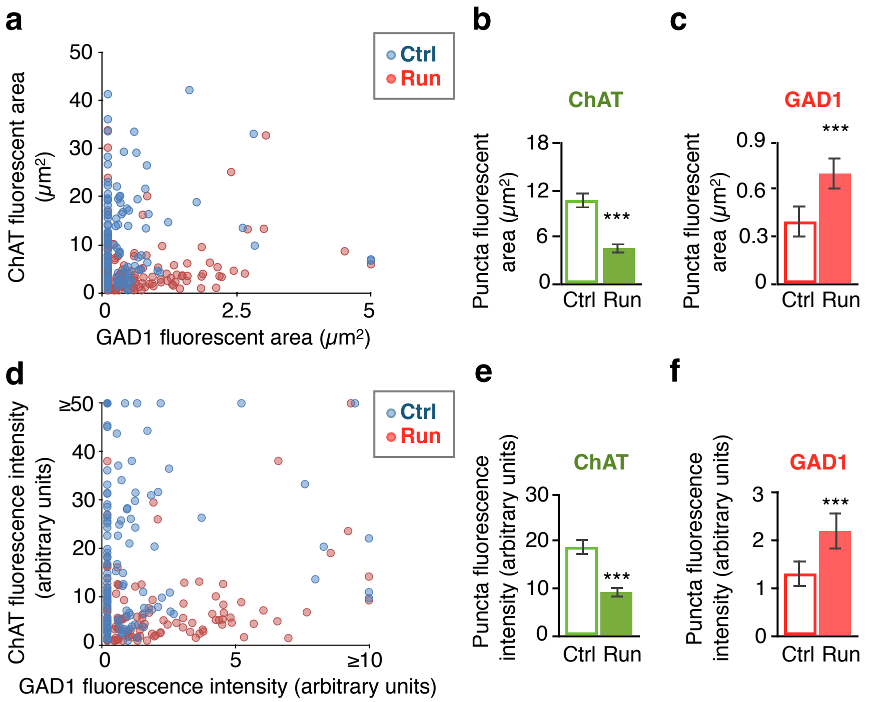


**Supplementary Figure 9**

**Changes of ChAT and GAD1 transcript area and intensity in nNOS neurons.**

**a**, Scatterplot of fluorescent area of ChAT transcripts (y-axis) against fluorescent area of GAD1 transcripts (x-axis) in nNOS neurons. Each dot represents one neuron. **b,c**, Mean fluorescent area of ChAT (**b**) or GAD1 (**c**) transcripts in single nNOS+ cells. **d**, Scatterplot of fluorescence intensity of ChAT transcripts (y-axis) against fluorescence intensity of GAD1 transcripts (x-axis) in nNOS cells. Each dot represents one neuron. **e,f**, Mean fluorescence intensity of ChAT (**e**) or GAD1 (**f**) transcripts in single nNOS+ neurons. For (**a**-**f**), n=4 animals/group; n=123 cells for Ctrl and 137 cells for Run. Statistical significance ***$p$<0.001 was assessed by Welch's t-test. Data shown are mean± SEM.

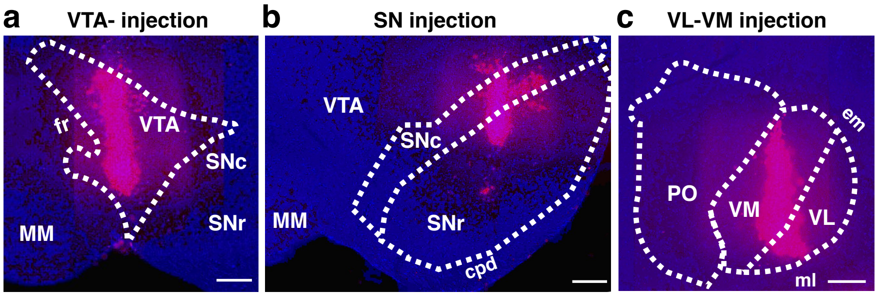


**Supplementary Figure 10**

**Anatomical evidence of retrobead injection.**

Representative coronal sections show retrobeads (magenta) injected into the VTA (**a**), SN (**b**) and VL-VM (**c**). MM, medial mammillary nucleus; fr, fasciculus retroflexus; SNc, substantia nigra, compact part; SNr, substantia nigra, reticular part; cpd, cerebral peduncle; PO, posterior complex of the thalamus; ml, medial lemniscus; em, external medullary lamina of the thalamus. The boundaries of the nuclei are drawn based on the comparison with the Allen mouse brain atlas. Scale bar, 200 µm.

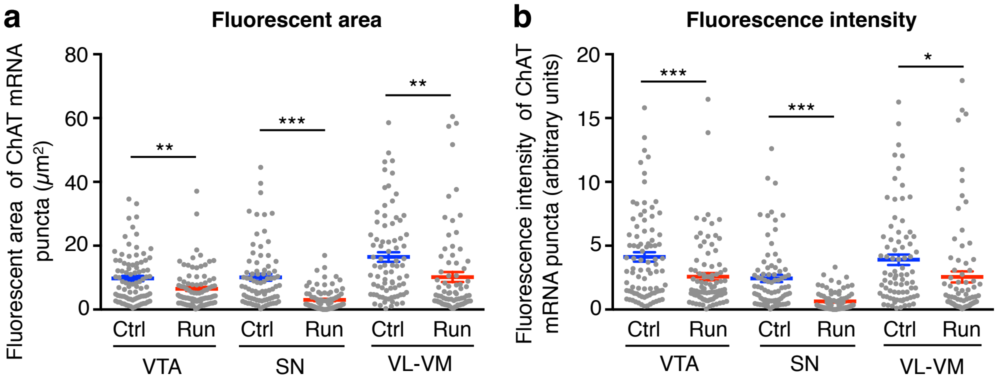


**Supplementary Figure 11**

**Fluorescent area and fluorescence intensity of ChAT transcripts in nNOS neurons that project to each brain region.**

Mean fluorescent area (**a**) or fluorescence intensity (**b**) of ChAT transcripts in nNOS neurons that project to corresponding brain regions (x-axis). Each dot represents one cell. n=4 animals/group. n=89 cells for VTA-Ctrl, 91 for VTA-Run, 81 for SN-Ctrl, 87 for SN-Run, 82 for VL-VM-Ctrl, 80 for VL-VM-Run. Statistical significance $^{*}p<0.05$, $^{**}p<0.01$, $^{***}p<0.001$ was assessed by Welch's t-test. Data shown are mean± SEM.

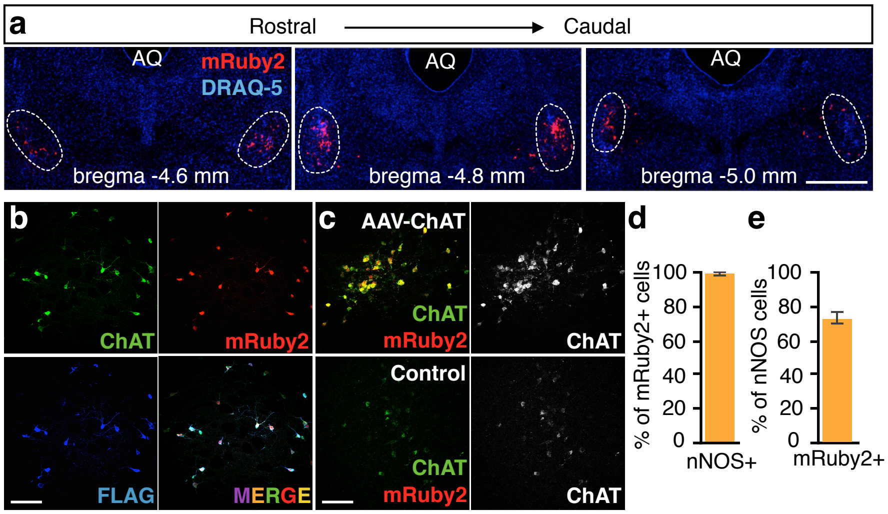


**Supplementary Figure 12**

**Validation of the use of the AAV-DIO-ChAT construct.**

**a,** mRuby2 expression in rostral-to-caudal sections of the cPPNs (dotted lines) of a single animal bilaterally-injected with AAV-DIO-ChAT. Coordinates adapted to Allen Brain Atlas. AQ, aqueduct. Scale bar, 1 mm. **b,** Triple-labeled images of ChAT, mRuby2, FLAG and merged image in a cPPN of an animal injected with AAV-DIO-ChAT. Scale bar, 100 µm. **c,** Double-stained images of ChAT and FLAG in both AAV-DIO-ChAT-injected (upper panels) or contralateral uninjected control (lower panels). Statistical analysis is in (**Fig. 5c**). Scale bar, 100 µm. **d**, The percentage of nNOS+mRuby2+ neurons in the mRuby2+ population (cell type specificity). n=4 animals/group. **e,** The percentage of nNOS+mRuby2+ population in the nNOS+ neurons (transfection efficiency). n=4 animals/group. Data shown are mean± SEM.

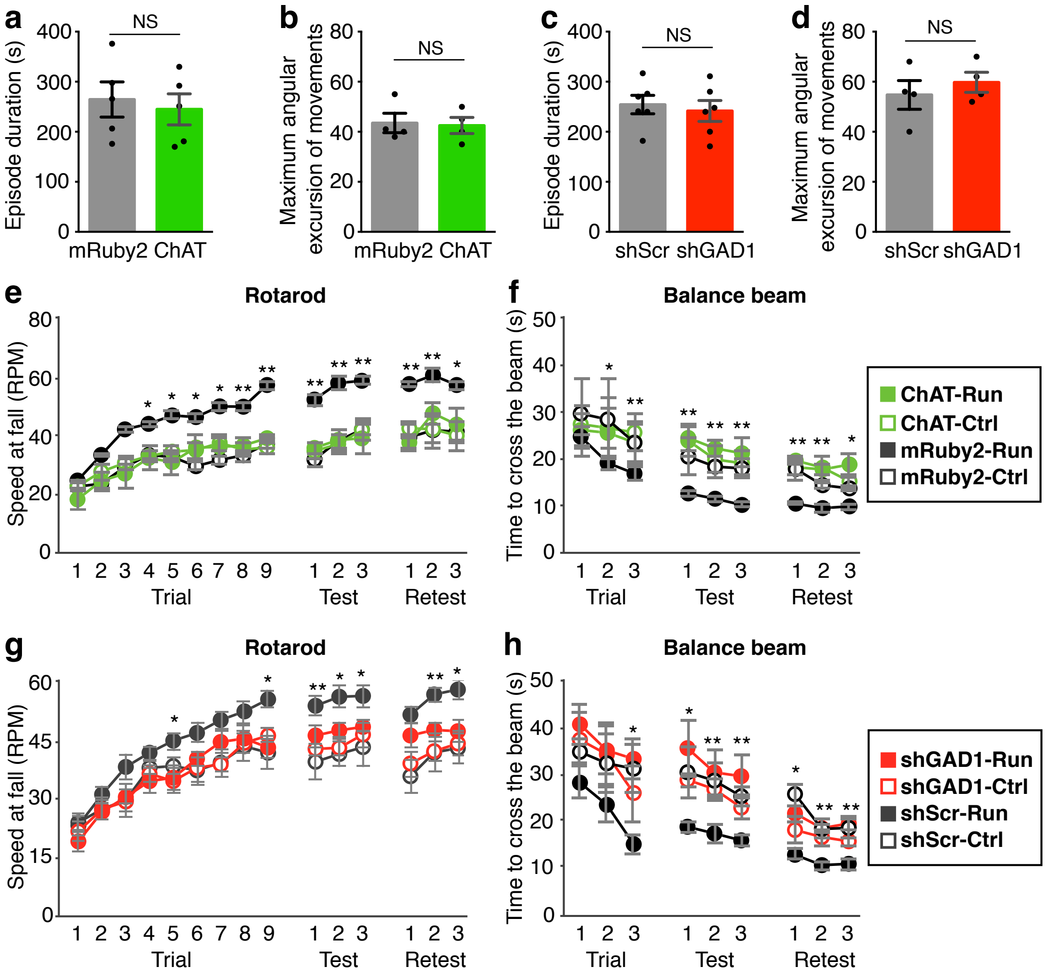


**Supplementary Figure 13**

**Running is not affected but running-induced improvements in motor skill learning are impaired when transmitter switching is overridden.**

**a-d,** Mean duration of running episodes and maximum angular excursion of mouse movements on the running wheels for 1-week trained ChAT-Cre runner mice that were injected with AAV-DIO-mRuby2, AAV-DIO-ChAT, AAV-DIO-shScr, or AAV-DIO-shGAD1 at the cPPN. For (**a**,**c**), n=5 animals for mRuby2 and ChAT, and 6 for shScr and shGAD1. For (**b**,**d**), n=4 animals for each group. **e**,**f**, Mean speed at fall on a rotarod and mean time to cross a 1 meter long, 0.75 meter high, 4 mm rod balance beam at each trial during training, at each test on the day after training, and at each retest 1 week after test. n=9 animals for mRuby2-Run and 18 animals for ChAT-Run. **g**,**h**, Same as (**e**,**f**). n=8 animals for shScr-Run and 9 for shGAD1-Run. Statistical significance *$p$<0.05, **$p$<0.01 was assessed by Welch's t-test. NS, not significant. Data shown are mean± SEM.

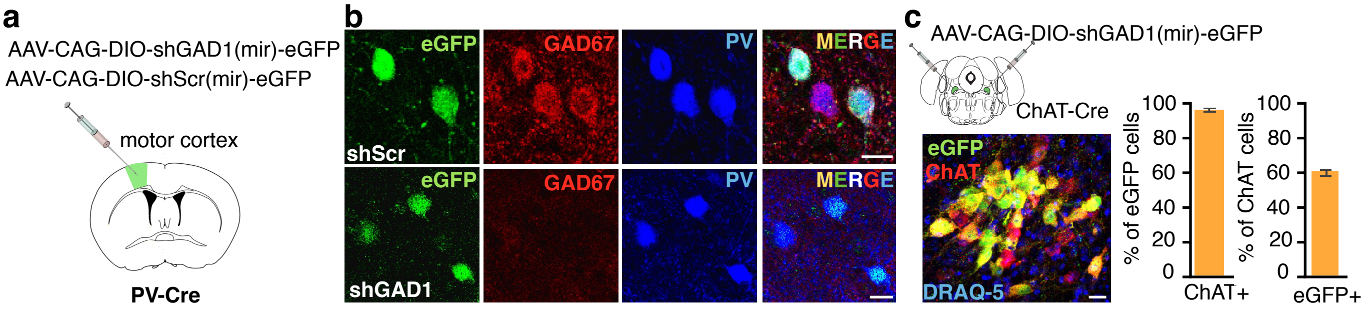


**Supplementary Figure 14**

**Validation of the use of the AAV-DIO-shGAD1 construct.**

**a,** Knockdown efficiency of GAD67 was verified *in vivo* by expressing either AAV-DIO-GAD1-shRNAmir-eGFP or AAV-DIO-scramble-shRNAmir-eGFP constructs in the cortex of a PV-Cre mouse. **b,** Brain sections from (**a**) were immunostained for GAD67 and PV**.** Scale bar, 20 µm. **c**, Triple-labeled images of eGFP, ChAT, and DRAQ-5 in the cPPN of a non-runner ChAT-IRES-Cre mouse injected with AAV-DIO-shGAD1. The specificity of transduction was measured as the percentage of ChAT+eGFP+ neurons in the eGFP population (>95%, 197/208) and the efficiency was measured as the percentage of ChAT+eGFP+ neurons in the ChAT population (~60%, 197/330). Similar specificity (>95%) and efficiency (~60%) were also observed for the control shScr construct. Scale bar, 20 µm. n=3 animals/group. Data shown are mean± SEM.

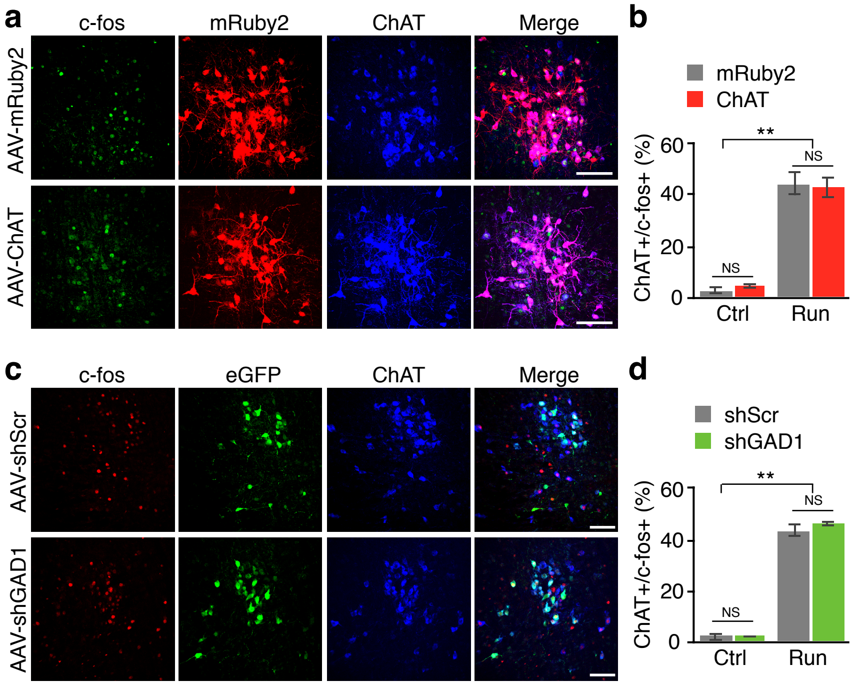


**Supplementary Figure 15**

**Overriding transmitter switching does not affect c-fos expression in cholinergic cPPN neurons.**

**a,** Triple labeling of c-fos, mRuby2 and ChAT in 1-week runners that were injected with AAV-DIO-mRuby2 or AAV-DIO-mRuby2-P2A-ChAT. Scale bar, 100 µm. **b,** Summary of (**a**) and non-runner controls. n=214 cells for mRuby2-Ctrl, 204 for ChAT-Ctrl, 223 for mRuby2-Run and 206 for ChAT-Run. n=3 animals/group. **c,** Triple labeling of c-fos, eGFP and ChAT in 1-week runners that were injected with AAV-DIO-shScr-eGFP or AAV-DIO-shGAD1-eGFP. Scale bar, 100 µm. **d,** Summary of (**c**) and non-runner controls. n=213 cells for shScr-Ctrl, 360 for shGAD1-Ctrl, 277 for shScr-Run and 315 for shGAD1-Run. n=3 animals/group. Statistical significance $^{**}p<0.01$ was assessed by ANOVA followed by Tukey's test. NS, not significant. Data shown are mean± SEM.
